# Mammalian SWI/SNF collaborates with a polycomb-associated protein to regulate male germ line transcription in the mouse

**DOI:** 10.1101/476143

**Authors:** Debashish U. Menon, Yoichiro Shibata, Weipeng Mu, Terry Magnuson

## Abstract

A deficiency in BRG1, the catalytic subunit of the SWI/SNF chromatin remodeling complex, results in a meiotic arrest during spermatogenesis. Here, we explore the causative mechanisms. BRG1 is preferentially enriched at active promoters of genes essential for spermatogonial pluripotency and meiosis. In contrast, BRG1 is also associated with the repression of somatic genes. Chromatin accessibility at these target promoters is dependent upon BRG1. These results favor a model where BRG1 coordinates spermatogenic transcription to ensure meiotic progression. In spermatocytes, BRG1 interacts with SCML2, a testes specific PRC1 factor that is associated with the repression of somatic genes. We present evidence to suggest that BRG1 and SCML2 concordantly regulate genes during meiosis. Furthermore, BRG1 is required for the proper localization of SCML2 and its associated deubiquitinase, USP7, to the sex chromosomes during pachynema. SCML2 associated, mono ubiquitination of histone H2A lysine 119 (H2AK119ub1) and acetylation of histone lysine 27 (H3K27ac) are elevated in *Brg1^cKO^* testes. Coincidentally, the PRC1 ubiquitin ligase, RNF2 is activated while a histone H2A/H2B deubiquitinase, USP3 is repressed. Thus, BRG1 impacts the male epigenome by influencing the localization and expression of epigenetic modifiers. This mechanism highlights a novel paradigm of co-operativity between SWI/SNF and PRC1.

**Summary statement:** BRG1, a catalytic subunit of SWI/SNF chromatin remodeling complex, interacts with SCML2 (<u>S</u>ex <u>c</u>omb on <u>m</u>idleg-like 2), a polycomb repressive 1 (PRC1) factor, to regulate transcription during spermatogenesis. This represents a novel paradigm of SWI/SNF-PRC1 co-operation during gametogenesis.

## Introduction

Spermatogenesis is a developmental cascade in which genetic information is passed on from mitotic precursors to meiotically derived haploid gametes. This process is particularly sensitive to the activity of several epigenetic regulators, known to influence meiotic recombination. Examples include the role of the meiosis specific H3K4 methyltransferase PR domain zinc finger protein9 (PRDM9) in double stranded break (DSB) formation, the roles of polycomb repressive complex2 (PRC2) and the H3K9 methyl transferases, EHMT2- Euchromatic histone lysine N-methyltransferase 2 (G9a); Suppressor of variegation 3-9 1 or 2 (SUV39H1/H2) in homolog pairing (synapsis) (Brick et al., 2012; Diagouraga et al., 2018; Hayashi et al., 2005; Mu et al., 2014; Tachibana et al., 2007; Takada et al., 2011). Other critical activities include the regulation of spermatogenic transcription by the polycomb repressive complexes, PRC1 and PRC2 (Hasegawa et al., 2015; Mu et al., 2014). Furthermore the ATP dependent family of nucleosome remodelers such as INO80, SWI/SNF and CHD5, known to modulate chromatin accessibility are also essential for spermatogenesis. (Kim et al., 2012; Li et al., 2014; Serber et al., 2015). In summary, chromatin modifiers play a vital role in germ line development.

Our lab previously reported the role of BRG1 (SMARCA4 – SWI/SNF catalytic subunit) in male meiosis (Kim et al., 2012). Briefly, the germ line depletion of BRG1 results in pachytene arrest. Mutant spermatocytes display unrepaired DNA DSBs evidenced by persistent γH2Ax, chromosomal asynapsis, and reduced MLH1 foci, a marker of crossovers (Kim et al., 2012, Wang et al. 2012). Coincidentally, an enhanced level of repressive chromatin is observed in mutant spermatocytes. Since SWI/SNF modulates chromatin accessibility by either sliding or evicting nucleosomes (Clapier et al., 2017), it is plausible that changes in chromatin structure seen in the *Brg1^cKO^* testes might result in meiotic defects by potentially influencing transcription or DNA repair. Both processes are influenced by SWI/SNF.

SWI/SNF is associated with both gene activation and repression. In mouse embryonic fibroblasts, BRG1 and the core SWI/SNF subunit, SNF5 co-ordinate nucleosome occupancy at promoters to achieve transcriptional regulation (Tolstorukov et al., 2013). The transcriptional outcome is often dictated by the subunit composition of the complex, which in turn can influence its genome-wide association and interactions with other gene regulators (Euskirchen et al., 2011; Raab et al., 2015; Raab et al., 2017). This is a theme that features frequently in the regulation of tissue and cell type specific transcriptional programs that impact critical processes such as embryonic stem (ES) cell pluripotency and differentiation, neuronal and cardiac cell fate specification (Ho and Crabtree, 2010). Thus, SWI/SNF directed gene regulation plays a critical role during development.

Apart from gene regulation, SWI/SNF has also been implicated in several DNA repair mechanisms. In Yeast and cell culture models, SWI/SNF is recruited to sites of DNA damage to promote accessibility and stimulate γH2Ax, a key component of the DNA damage response (DDR) signaling pathway (Kwon et al., 2015; Lee et al., 2010; Ogiwara et al., 2011). Other activities include the recruitment of homologous recombination (HR) and non-homologous end joining (NHEJ) repair factors (Ogiwara et al., 2011; Qi et al., 2014; Watanabe et al., 2014), the transcriptional silencing of regions adjacent to DNA DSB (Kakarougkas et al., 2013). While BRG1 is dispensable for γH2Ax formation during meiosis, it has been reported to influence the distribution of DDR factors like RAD51 (DNA recombinase) and RPA (Replication protein A1) (Kim et al., 2012).

In this study we present evidence to show that BRG1 coordinates spermatogenic transcription. BRG1 activates genes essential for maintaining spermatogonial pluripotency and meiotic progression. In contrast it represses somatic genes. Our data suggest that somatic gene repression is achieved through an interaction with SCML2 (<u>S</u>ex <u>c</u>omb on <u>m</u>idleg-like 2), a known testis specific PRC1 (Polycomb Repressive Complex 1) member (Luo et al., 2015). BRG1 is required for the normal localization of SCML2 and its interacting partner, USP7 (Ubiquitin Specific Peptidase 7) during pachynema, suggesting a role in recruitment. Furthermore, histone modifications associated with SCML2, such as the repressive mono ubiquitination of histone H2A lysine 119 (H2AK119ub1) and activating acetylation of histone lysine 27 (H3K27ac) are perturbed in *Brg1^cKO^* testes. Coincidentally, BRG1 also activates the expression of the H2A ubiquitin ligase, RNF2 (Ring Finger Protein 2) and represses the histone H2A/H2B deubiquitinase, USP3 (Ubiquitin Specific Peptidase 3). Thus SWI/SNF can epigenetically regulate germ line transcription by SCML2 dependent and independent mechanisms.

## Results

### BRG1 associates with transcriptionally active and poised chromatin

To understand the functions of SWI/SNF activity during meiosis, we determined the genome-wide association of BRG1 by ChIP-seq. We reasoned that the examination of BRG1 occupancy concurrent to critical meiotic processes such as DSB repair and homologous chromosome synapsis would yield insight into the molecular mechanisms underlying the meiotic defects observed in the *Brg1*^*fl/Δ*^; *Mvh-cre*^*Tg/0*^ (*Brg1^cKO^*; see materials and methods) males. For this purpose we isolated chromatin from spermatogenic cells obtained from P12 (mostly pre-pachytene germ cells) and P18 testes (predominantly pachytene spermatocytes) (Bellve et al., 1977; Goetz et al., 1984). At these stages, BRG1 appears to be enriched promoter proximally (Fig. 1A, panels 1 & 4). This is consistent with the fact that more than 50% of BRG1 peaks associate with promoters at P12 and P18, while only a minority map to distal sites (Fig. S1A). By performing K-means clustering we categorized transcription start sites (TSS) into three different classes (class1: Cl-1, class2: Cl-2, class: Cl-3) based upon their association with BRG1. These TSS’s feature high (Cl-1), medium (Cl-2) and insignificant (Cl-3) BRG1 enrichment.

**Figure 1.**
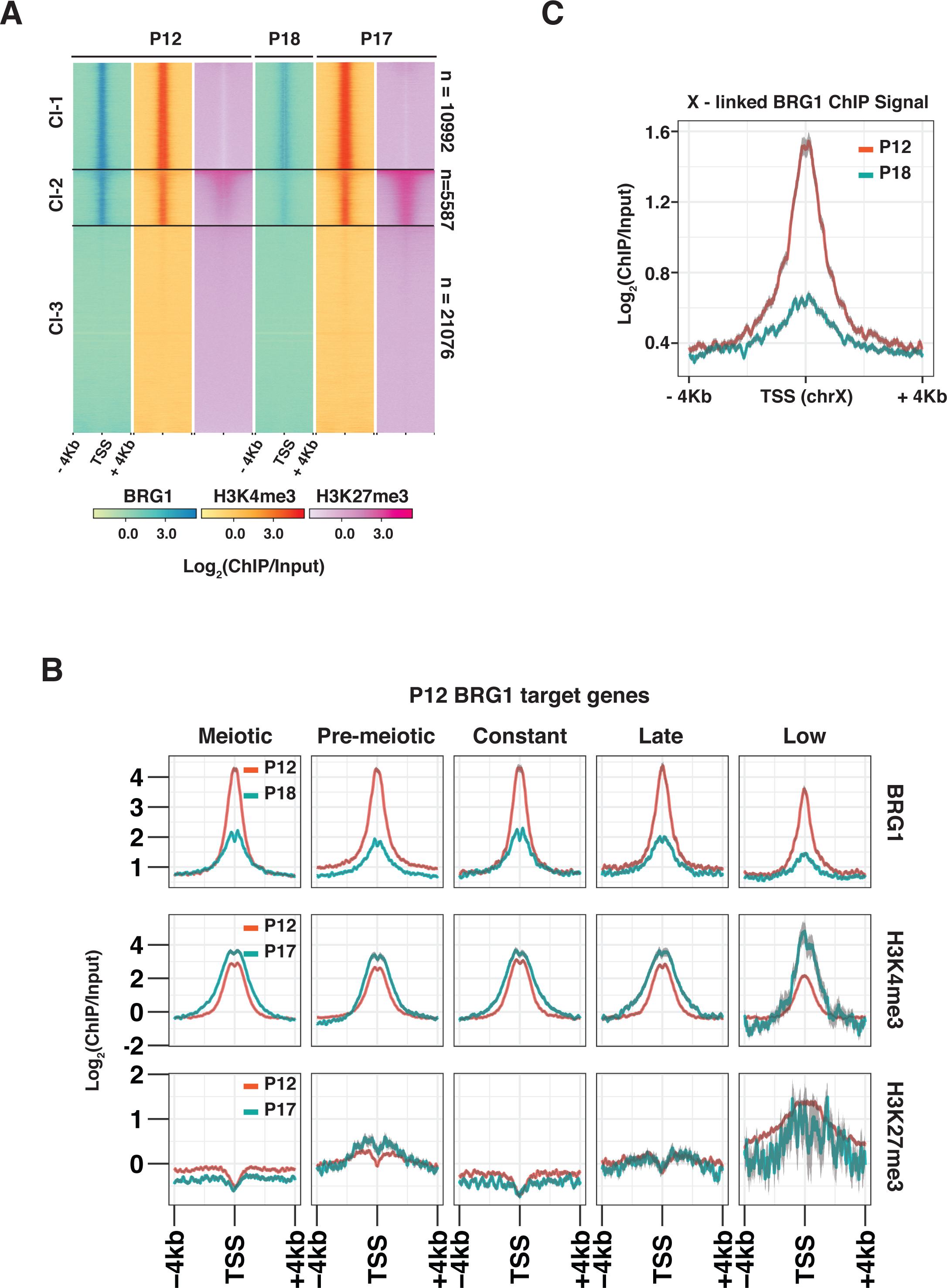
BRG1 is enriched at transcriptionally active and poised regions. The relative enrichment of BRG1, H3K4me3 and H3K27me3 from P12, P17 and P18 testes at, (A) RefSeq gene, TSS ± 4 Kb shown using heatmap with K-means clustering (B) TSS ± 4 Kb associated with BRG1 target (P12 peaks) genes categorized by their temporal expression profile. (C) P12 and P18 BRG1 enrichment at TSS ± 4 Kb on chromosome X. TSS: Transcription start site.

To gain insight into the activity of genes associated with BRG1 in spermatogenic cells, we surveyed the chromatin environment surrounding their TSS’s. We monitored the enrichment of activating trimethylation of histone H3 lysine 4 (H3K4me3) and repressive trimethylation of histone H3 lysine 27 (H3K27me3), associated with class1-3 TSS’s at P12 and P17 (Mu et al., 2014) (Fig. 1A). Robust H3K4me3 enrichment appeared to be associated only with Cl-1 and Cl-2 TSS’s from P12 to P17 (Fig. 1A, panels 2 & 5). In contrast, H3K27me3 levels at P12 and P17 appear depleted at Cl-1 TSS’s featuring high BRG1 occupancy (Fig. 1A, panels 3 & 6). Such antagonism is a well-established feature of SWI/SNF and PRC2 genomic associations (Wilson et al., 2010). Cl-3 TSS’s only displayed basal levels of H3K27me3, which is particularly discernable at P17. Unlike Cl-1 and Cl-3, Cl-2 TSS’s display significantly higher levels of H3K27me3, which appears progressively enhanced from P12 to P17 (Fig. 1A, panels 3 & 6). The co-occurrence of H3K27me3 and H3K4me3 at Cl-2 TSS’s resemble features of bivalent promoters, which are usually associated with transcriptionally quiescent genes poised to be reactivated at later stages of development (Bernstein et al., 2006; Hammoud et al., 2014; Lesch et al.,2013). Thus genes associated with Cl-2 TSS’s likely represent repressed BRG1 target genes. In contrast potential activated gene targets appear to be associated with Cl-1 TSS’s.

To understand BRG1 directed gene regulation in the context of spermatogenesis, we categorized genes associated with each K-means cluster (Fig. 1A), by their temporal expression profiles, previously determined from whole-testes RNA seq data (Margolin et al., 2014). These include genes maximally expressed in testes, at P6 (pre-meiotic: Pmei), from P8-P20 (meiotic: Mei), at P38 (Late, adult testis), from P6-P38 (Constantly; Const) and those expressed at low levels in testes from P6-P38 (Low, < 2 Reads per Kilobase per Million of Reads; RPKM). The majority of Cl-1 associated genes were meiotic with relatively fewer pre-meiotic genes. In contrast, Cl-2 and Cl-3 were mostly comprised of pre-meiotic and low genes (Fig. S1B). This is particularly interesting given that Cl-1 TSS’s are exclusively marked by H3K4me3, while Cl-2 promoters display bivalent chromatin modifications. We observed similar trends when monitoring H3K4me3 and H3K27me3 dynamics, at TSS’s of BRG1 target genes (P12 peaks) categorized by their temporal expression profile during meiosis (P12 to P17) (Fig. 1B). Over this duration all gene categories experience a 4-fold decrease in BRG1 enrichment in the presence of abundant H3K4me3 (Fig. 1B, top and middle row). Only pre-meiotic and low gene targets display elevated levels of H3K27me3 from P12 to P17 (Fig. 1B, bottom row), distinguishing them from, meiotic, constant and late gene targets, that appear exclusively marked by H3K4me3 (Fig. 1B, middle and bottom row). Candidate pre-meiotic targets such as *Zbtb16* and *Id4*, which are markers of undifferentiated spermatogonial cells along with *Pdgfra*, a somatic signaling receptor, that are normally repressed during meiosis, display bivalent promoters (Basciani et al., 2002; Green et al., 2018; Hammoud et al., 2014). In contrast, meiotic target, *Sycp1*, which is essential for synaptonemal complex assembly displays a H3K4me3 enriched promoter (Fig. S1C) (De Vries et al., 2005). Thus BRG1 might co-ordinate the expression of genes over the measured course of spermatogenesis.

Given its association with active (H3K4me3) and poised (H3K4me3/H3K37me3) chromatin, we were curious to examine BRG1 localization to the sex chromosomes, which are transcriptionally silenced during pachynema (reviewed in Turner, 2007). We examined BRG1 occupancy at the TSS’s of X – linked genes at P12 (Pre-pachytene stages) and P18 (Pachytene stages). BRG1 enrichment at X – linked TSS’s appeared reduced at P18 relative to P12, but not absent (Fig. 1C). Thus BRG1 associates with meiotically inactivated sex chromosome.

Apart from TSS’s, H3K4me3 is also enriched at DSB/recombination hotspots, known to be associated with a meiosis specific histone methyl transferase, PRDM9 (Brick et al., 2012; Diagouraga et al., 2018; Hayashi et al., 2005). We therefore examined BRG1 association at these PRDM9 sites previously mapped by ChIP-seq in P12 testes (Baker et al., 2015). The lack of enrichment at PRDM9 peaks makes it unlikely that BRG1 directly affects DSB formation (Fig. S1D). Thus BRG1 might play a major role in gene regulation during meiosis.

### BRG1 co-ordinates spermatogenic gene expression

The promoter centric association of BRG1 prompted us to examine its influence on the transcription of target genes by RNA-seq. We compared transcript abundance between spermatogenic cells isolated from P12 *Brg1^fl/+^* (*Brg1^WT^*) and *Brg1*^*fl/Δ*^; *Mvh-cre*^*Tg/0*^ (*Brg1*^*cKO*^) testes, where the germ cell populations are mostly pre-pachytene and therefore unlikely to be influenced by pachytene arrest. In fact, the loss of BRG1 did not appear to affect the development of pre-pachytene spermatocytes as staged by γH2Ax (meiotic marker) at P10 and P13 (Fig. S2A, left panel). Incidence of pachytene arrest only manifest at P14 (Fig. S2A, left panel). In agreement with these results, the abundance of pre-pachytene protein coding transcripts (Ball et al., 2016) appeared similar between P12 *Brg1*^*WT*^ and *Brg1*^*cKO*^ testes (Fig. S2A). Only, early and late pachytene specific transcripts were slightly less abundant upon the loss of BRG1 at P12, predictive of pachytene arrest at later stages (Fig. S2A, right panel). Overall we do not expect the analysis of gene expression to be impacted significantly by secondary effects such as developmental delays.

To identify genes significantly mis-expressed (FDR ≤ 0.05) upon the loss of BRG1, we performed an edgeR analysis on the RNA-seq data. An equivalent number of genes are either transcriptionally down regulated (n=1100) or up regulated (n=983) in P12 *Brg1^cKO^* relative to *Brg1^WT^* testes (Fig. 2A). More genes were down regulated (n=310) by a magnitude of 2-fold or higher, relative to those up regulated (n=75), upon the loss of BRG1 (Fig. 2A). Nearly half of these differentially regulated genes were associated with BRG1 peaks (P12 peaks). The down regulated genes appear normally expressed in gonadal tissue and the nervous system and were enriched for Gene ontology (GO) terms relevant to meiotic processes (Fig. 2B). In contrast, the up regulated genes are normally expressed in limb and muscle and were enriched for GO terms relevant to somatic developmental processes (Fig. 2B). Thus, BRG1 coordinates germ line transcription by activating meiotic genes while concomitantly repressing somatic genes. Since BRG1 is essential for meiotic progression (Kim et al., 2012), we monitored the expression of genes associated with abnormal spermatogonia proliferation and meiotic arrest phenotypes, curated from the mouse genome database (Blake et al., 2003). We generated a heatmap displaying z-scores associated with transcript abundance of each candidate gene, measured across P12 *Brg1^WT^* and *Brg1^cKO^* replicates (Fig. 2C). With the exception of *Androgen receptor*(*Ar*) and *Microtubule associated protein 7* (*Map7*), all other germ cell factors appeared less abundant upon the loss of BRG1 (Fig. 2C). Next, we adopted a reverse genetic approach to test whether the mis-expression of candidate genes were associated with specific phenotypes in the *Brg1^cKO^*. To do this we choose a pre-meiotic candidate, *Zbtb16* and meiotic candidate, *Sycp2*.

**Figure 2.**
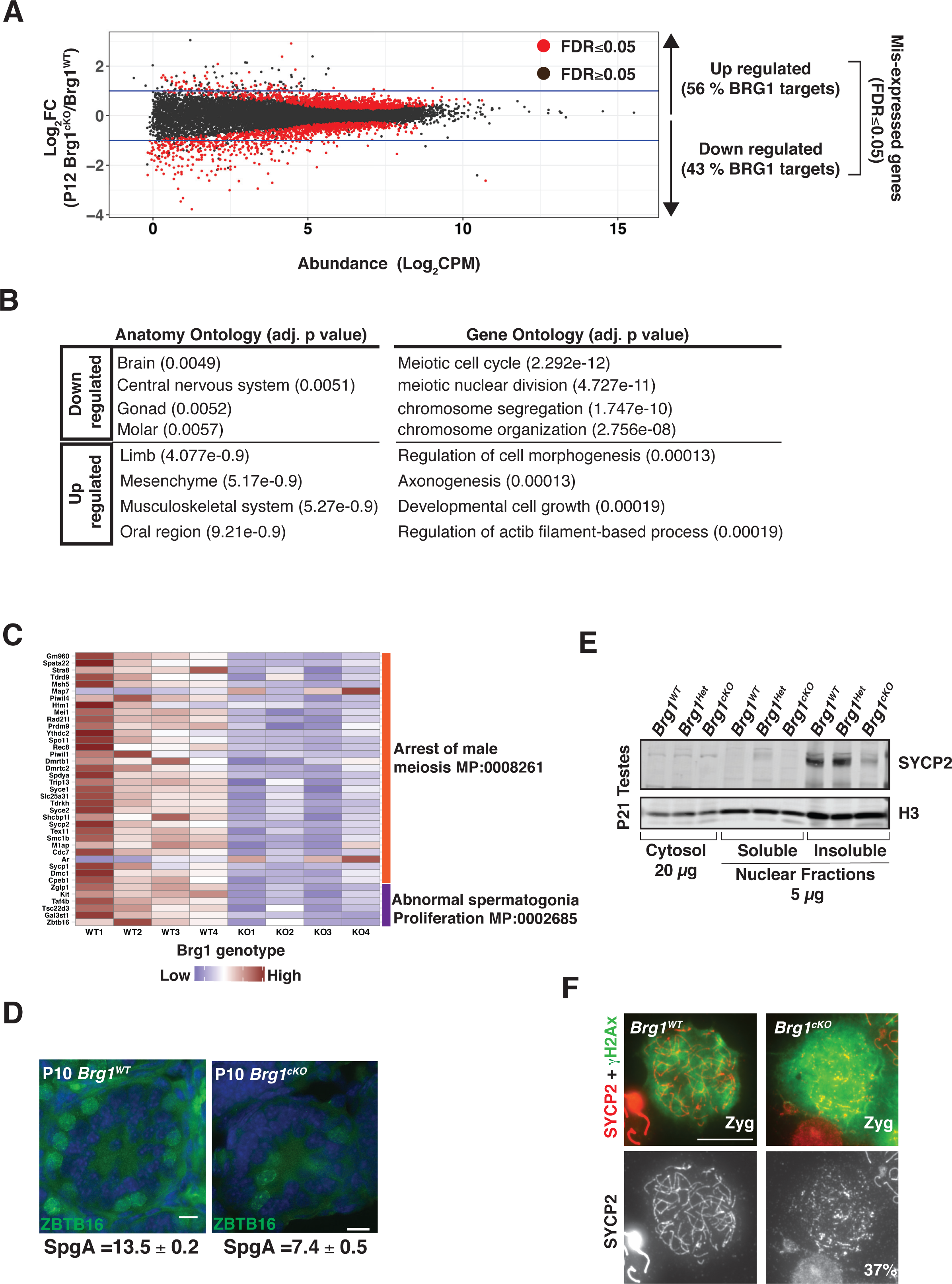
BRG1 influences transcription during spermatogenesis. (A) Log2 fold change (y-axis) in transcript abundance (CPM: Counts per million, x-axis) of genes in the P12 *Brg1^cKO^* relative to *Brg1^WT^*. Each dot denotes a gene and horizontal blue lines denote a 2-fold. (B) Table listing enriched anatomy and gene ontology terms associated with BRG1 regulated genes. Benjamini – Hochberg, adjusted p-values are reported in parenthesis. (C) Heatmap showing transcript abundance (z-scores) of genes (rows) associated with mouse phenotype ontologies in P12 *Brg^WT^* (WT1-WT4) and *Brg^cKO^* (KO1-KO4) (columns). (D) P10 *Brg^WT^* and *Brg1^cKO^* testes cryosections (25x objective Scale bar: 20 μm), immuno-labeled for ZBTB16 (green) and counter stained with DAPI (blue). The average numbers of SpgA and standard error of measurement (SEM) are indicated. (E) Western blot for SYCP2 in sub-cellular fractions obtained from P21 *Brg1^WT^*, *Brg^Het^* and *Brg1^cKO^* spermatogenic cells. Histone H3 is loading control. (F) *Brg^WT^* and *Brg1^cKO^* zygotene spermatocytes (100x objective, Scale bar: 10 μm) immuno-labeled for SYCP2 (red) and γH2Ax (green).

*Zbtb16* is essential for the maintenance of a pool of undifferentiated typeA spermatogonia (SpgA) (Buaas et al., 2004). While BRG1 is dispensable for the establishment of spermatogonia, its role in SpgA maintenance remains unknown (Kim et al., 2012). We quantified the number of SpgA in P10 *Brg1^WT^* and *Brg1^cKO^* testes cryosections by immunostaining for ZBTB16. The *Brg1^cKO^* testes contain 46% fewer SpgA, when compared to *Brg1^WT^* testes (Fig. 2D). To validate the transcriptional basis of this defect, we performed qRT-PCR to determine the expression of *Zbtb16* in purified, THY1^+^ spermatogonia (enriched for spgA) isolated from P8 *Brg1^WT^*, *Brg1^Het^* (*Brg1*^*fl/+*^;*Mvh-cre*^*Tg/0*^) and *Brg1*^*cKO*^ testes. We also monitored the expression of other established stem cell factors such as, *Inhibitor of DNA binding 4* (*Id4*), *POU domain, class 5, transcription factor 1* (*Pou5f1*, also known as *Oct4*), *POU domain, class 3, transcription factor 1* (*Pou3f1, also known as Oct6*) and *Forkhead box O1* (*Foxo1*) (Dann et al., 2008; Goertz et al., 2011; Oatley et al., 2011; Wu et al., 2010). Transcripts associated with *Zbtb16*, *Id4* and *Pou5f1* were down-regulated upon the loss of BRG1 in the purified THY1^+^ fractions (Fig. S2B). The fact that *Thy1* mRNA levels remain abundant argues against a loss of spermatogonia cells early in development (Fig. S2B). Thus, BRG1 regulates the maintenance of undifferentiated spermatogonial cells by activating the expression of critical stem cell factors.

The meiotic gene candidate, *Sycp2*, constitutes the structural component of the lateral element of meiotic chromosomal axes and is essential for synapsis (Yang et al., 2006). Coincidentally, *Brg1^cKO^* spermatocytes display an increase in asynapsis (Kim et al., 2012). In the *Brg1^cKO^*, SCYP2 levels are distinctly lower relative to the controls (Fig. 2E, Fig. S2C) and abnormally assemble into short lateral filaments and aggregates in 37% of the mutant zygotene spermatocytes (total scored=212) (Fig. 2F, Fig. S2D). While *Sycp2^-/-^* spermatocytes fail to form SYCP3 elements, we did not observe a similar defect in the *Brg1^cKO^* spermatocytes (Fig. S2D) (Yang et al., 2006). We reason that the reduced levels of SYCP2 might be sufficient to facilitate apparently normal SYCP3 assembly. Thus, a paucity in SYCP2 may limit meiotic progression by potentially compromising the formation of a functional synaptonemal complex. In addition to the mis-regulation of essential germ cell factors, we also identified a significant increase in the expression of several somatic genes in the *Brg1^cKO^* (Table S1). *Pdgfra* a signaling receptor, generally associated with somatic (Basciani et al., 2018; Schmahl et al., 2008) and pre-meiotic spermatogonia (Hammoud et al., 2014), was up-regulated by more than 2-fold over a period spanning meiosis-I in the *Brg1^cKO^* (Fig. S2E).

### BRG1 is required to maintain chromatin accessibility at promoters

Chromatin remodelers reposition nucleosomes, consequently regulating accessibility to transcription factors. Thus BRG1 might influence transcription by modulating the structure of the germ line epigenome. To investigate this, we performed Assay for Transposase-Accessible Chromatin (ATAC)-seq to map open chromatin in pre-pachytene and pachytene spermatogenic cells isolated from P12 and P18 testes respectively. Similar to the RNA-seq data, we surveyed differences in chromatin accessibility between P12 *Brg1^WT^* and *Brg1^cKO^* testes (Fig. S3A, left panel). Since P18 *Brg1^cKO^* testes are characterized by severe pachytene arrest, we compared chromatin accessibility between P18 *Brg1^WT^* and *Brg1^Het^* (*Brg1^fl/Δ^*) testes (Fig. S3A, right panel). In normal spermatogenic cells, strong ATAC read coverage was detected promoter proximally at both P12 and P18 (Fig. S3A). This promoter accessibility is significantly diminished in P12 *Brg1^cKO^* testes and also under conditions of haploinsufficiency in P18 *Brg1^Het^* testes (FigS3A).

Since promoter proximal chromatin responds to the loss of BRG1, we first monitored the changes in chromatin accessibility at promoters of target genes differentially regulated by BRG1 (Fig. 3A). Promoters of BRG1 target genes that are normally activated (down regulated upon BRG1 loss), display a significant decrease in promoter accessibility in the *Brg1^cKO^* relative to *Brg1^WT^* testes. While the promoters of repressed gene targets (up regulated upon BRG1 loss) are less accessible relative to their activated counterparts in *Brg1^WT^* testes, their accessibility appears further reduced in the *Brg1^cKO^* testes, albeit at levels that fail to meet statistical significance (Fig. 3A). Thus only activated gene promoters experience a significant change in accessibility.

**Figure 3.**
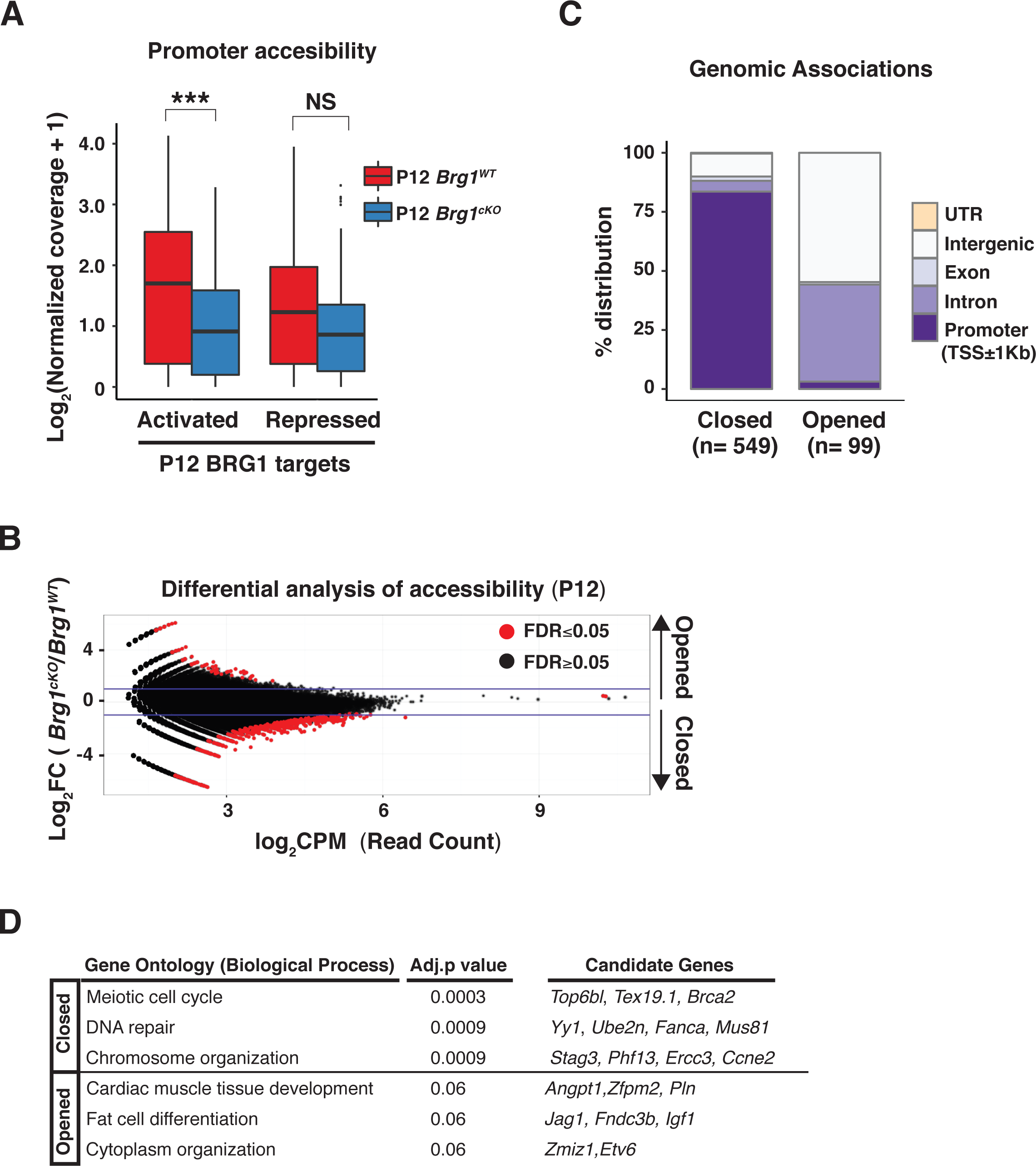
BRG1 directly regulates chromatin accessibility at promoters. (A) Log2 normalized ATAC-seq read coverage with added pseudocount (y-axis) at promoters (TSS± 0.5Kb) of BRG1 regulated genes (x-axis) in P12 *Brg1^wt^* (red) and *Brg1^cKO^* (blue) spermatogenic cells. ***: p<0.001; NS: not significant, as calculated by Wilcoxon rank sum test (B) Log2 fold change (y-axis) in read counts (CPM: counts per million, x-axis) in P12 *Brg1^cKO^* relative to *Brg1^WT^*. Each dot represents a 300bp-binned region. Horizontal blue lines denote 2-fold change. (C) Genomic associations of closed and opened regions. n: Total number of regions. (D) Gene ontology analysis of closed and opened regions. Benjamini – Hochberg, adjusted p-values are shown.

To identify genome-wide changes in chromatin accessibility at P12, we performed edgeR on the ATAC read counts obtained from wild type and mutant samples. The vast majority of regions that displayed significant differences in chromatin accessibility appear less accessible (Closed; n= 549), leaving only a few regions that acquire greater accessibility (Opened; n= 99) upon the loss of BRG1 at P12 (Fig. 3B). Consistent with the general decrease in promoter accessibility in the *Brg1^cKO^* (Fig. S3A), the closed regions are overwhelmingly associated with promoters (Fig. 3C, Fig. S3B; panel 1). In contrast, the opened regions were prominently featured within introns and intergenic regions (Fig. 3C, Fig. S3B, panel 2 & 3). This overall decrease in chromatin accessibility genome-wide is consistent with the previously observed increase in repressive epigenetic modifications in *Brg1^cKO^* spermatocytes (Kim et al., 2012). The genes associated with closed promoters in *Brg1^cKO^* testes were mostly meiotic in function (Fig. 3D, Table S2). Thus BRG1 probably activates meiotic genes by maintaining chromatin accessibility at cognate promoters. In contrast a few genes associated with the distal sites that appear more accessible in the *Brg1^cKO^* testes, represent somatic factors (Fig. 3D, Table S2).

### BRG1 physically interacts with SCML2, a non-canonical PRC1 factor

To investigate further mechanisms governing SWI/SNF mediated epigenetic regulation, we monitored BRG1 interactions in testes nuclear extracts from 3-week-old mice by performing Immuno-pulldowns (IP) (Fig. 4A). Proteins isolated from a BRG1 IP and control nonspecific (ns) IgG pulldown were identified by mass spectrometry (MS). Known SWI/SNF sub-units were specifically identified in the BRG1 IP, thus demonstrating the efficacy of our method (Fig. 4A, Fig. S4A). Furthermore the presence of both SWI/SNF sub complexes; *Brahma* Associated Factor (BAF) and *polybromo*-BAF (PBAF) are detected in the germ line (Fig. 4A,Fig. S4A) (reviwed in Masliah-Planchon et al., 2015). More interestingly we identified peptides associated with SCML2, a known testes specific PRC1 factor (Hasegawa et al., 2015; Luo et al., 2015) (Fig. 4A). Since candidate peptides were also detected in the non-specific IgG IP, we validated these interactions by performing a reverse IP with an antibody against SCML2 (Fig. 4B). The IP was conducted on nuclear lysates treated with universal nuclease (benzonase) to eliminate non-specific, DNA mediated interactions. BRG1 was specifically detected in the SCML2 IP, compared to nsIgG (Fig. 4B, lane 2 & 3). Additionally, a previously established interaction between SCML2 and RNF2 was validated (Fig. 4B, lane3) (Hasegawa et al., 2015). A smear like appearance of SCML2 in the nuclear extracts (Fig 4B) prompted us to confirm the specificity of the SCML2 antibody. We did this by performing western blots on nuclear extracts obtained from spermatogenic cells and ovaries. Consistent with its male specific expression pattern, we fail to detect a SCML2 signal in nuclear extracts obtained from ovaries (Fig.S4B, lane 2). Hence the smearing might be a product of protein instability or indicative of isoforms.

**Figure 4.**
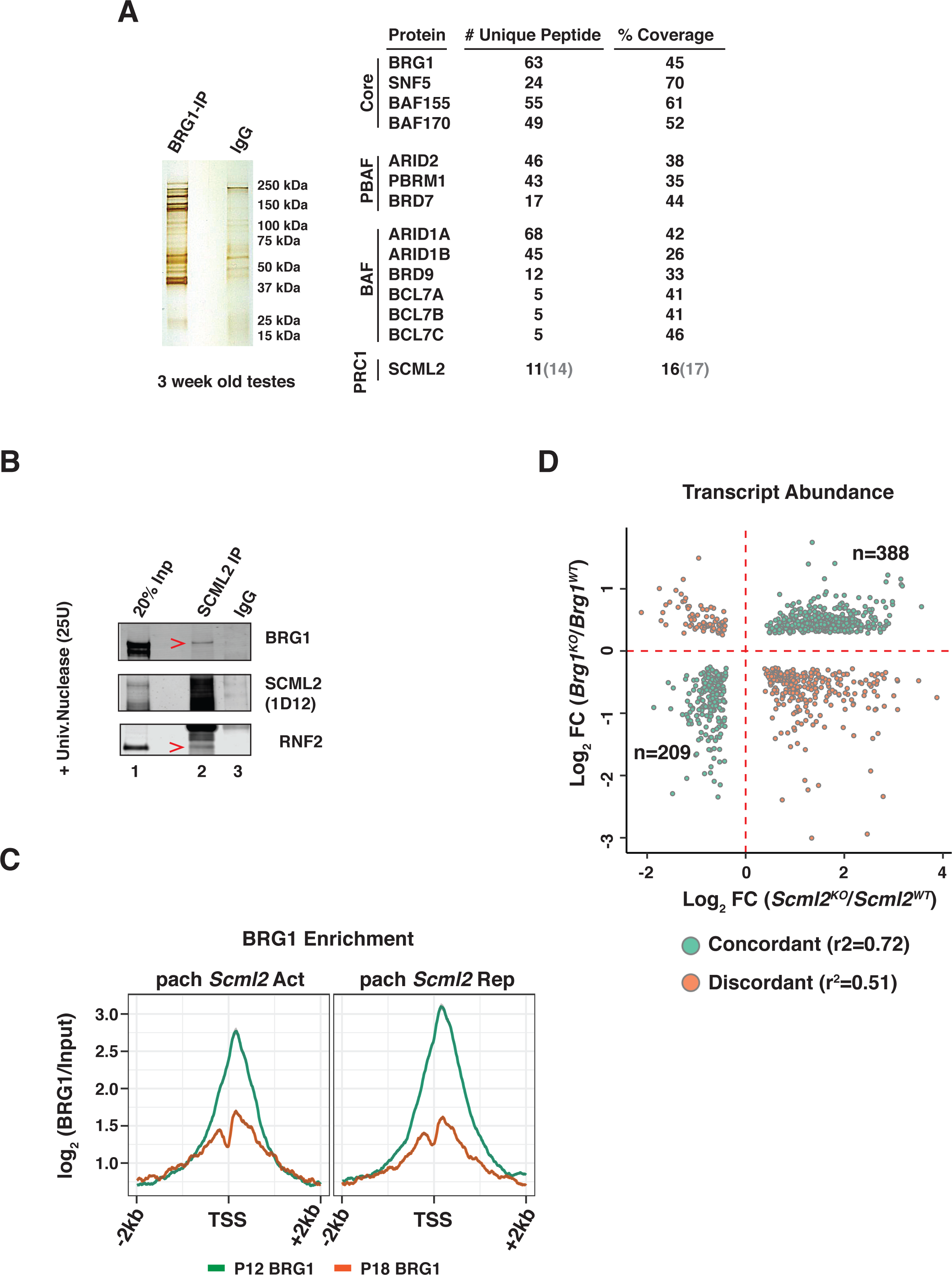
BRG1 physically interacts with SCML2 to regulate gene expression. (A) Silver stained gel (left) and table (right) summarizing BRG1 IP-MS results. Numbers in parenthesis are associated with control IgG. (B) SCML2 Co-IP analysis. Red arrowheads label interacting proteins. Lane numbers are labeled (1-3) (C) P12 (green line) and P18 (orange line) BRG1 enrichment at TSS ± 2 Kb, of genes differentially regulated by SCML2 in pachytene spermatocytes. pach *Scml2* Act: Activated, pach *Scml2* Rep: Repressed. (D) Log2 fold change in expression (KO/WT), of genes upon the loss of BRG1 (y-axis) and SCML2 (x-axis). Each dot represents a commonly mis-regulated gene (FDR<0.05), showing either concordant (green dots) or discordant (orange dots) changes in genes expression. n: number of concordantly mis-regulated genes. *r^2^* values were derived from pearson’s correlation test.

In pachytene spermatocytes, SCML2 is also known to interact with USP7, a known deubiquitinase and non-canonical member of the mammalian PRC1.4 complex (Lecona et al., 2015; Luo et al., 2015). Despite this association, USP7 does not interact directly with BRG1 (Fig. S4C, lane 2-4). Thus BRG1 only associates with SCML2 during meiosis.

### BRG1 and SCML2 concordantly regulate genes during meiosis

In pachytene spermatocytes, SCML2 is known to repress somatic and spermatogonial genes, while concurrently activating certain meiotic and late spermatogenic genes (Hasegawa et al., 2015). This is similar to the epigenetic role of BRG1 in the germ line. Hence it might be possible that BRG1 interacts with SCML2 to mutually regulate the germ line transcriptome. In support of such a possibility, we identified robust enrichment of BRG1 at TSS’s of genes differentially regulated (FDR < 0.05) by SCML2, during pachynema (Fig. 4C). The relative enrichment was greater at P12 (8 fold over input) compared to P18 (6 fold over input), indicating that BRG1 localizes prior to pachytene at this sites. Furthermore, chromatin accessibility at TSS’s associated with SCML2 regulated genes is reduced upon the loss of BRG1 at P12 (Fig. S5A).

Next, we looked for commonly mis-expressed genes (FDR<0.05) between the P12 BRG1 and pachytene SCML2 RNA-seq data. About 65% (n=597) of genes commonly mis-expressed in the absence of either BRG1 or SCML2 exhibited concordant expression changes that were highly correlated (r^2^=0.72). The remaining 35% (n=330) showed discordant changes (r^2^=0.51) (Fig. 4D). Of the concordantly regulated genes, 64% (n=388) were repressed, while the remainder (n=209) were activated. The concordantly repressed genes accounted for 40% of the genes up-regulated in the P12 *Brg1^cKO^* testes. In contrast, only 19% of genes down regulated in the P12 *Brg1^cKO^* testes overlapped with concordantly activated genes. The commonly repressed genes were mostly somatic in function (Fig. S5B) and included the somatic signaling receptor, PDGFRA, associated with BRG1 (Fig. S1C, Fig. S2E). Thus during normal prophase-I, BRG1 might achieve gene repression by recruiting SCML2. On the other hand the co-activated genes were associated with GO terms relevant to the mitotic spindle checkpoint (Fig S5B). Evidence for such checkpoint mechanisms have been reported late in meiosis (Eaker et al., 2001; Lee et al., 2011).

### BRG1 influences SCML2 and USP7 localization to the sex body

During pachynema, both SCML2 and its interacting partner, USP7 paint the sex body, a γH2Ax enriched, sub-nuclear compartment containing the sex linked chromosomes (Hasegawa et al., 2015; Luo et al., 2015). Therefore we determined whether SCML2 localization to the sex body was dependent on BRG1.

We first compared SCML2 localization in Brg1^WT^ and Brg1^cKO^ testes cryo-sections from 2- and 3-week-old males by IF (Fig. 5A,Fig. S6A). Mutant pachytene spermatocytes were identified by staining for γH2Ax, given that its association with the sex body remains unperturbed in the *Brg1^cKO^* (Kim et al., 2012). The loss of BRG1 appeared to impact the localization of SCML2 in *Brg1^cKO^* pachytene spermatocytes (Fig. 5A). Here, SCML2 appeared abnormally distributed genome-wide, without normally accumulating on the sex body (Fig. 5A, panel 3 insets). We confirmed these defects, by co-staining for ATR (Ataxia Telangiectasia and Rad3 Related), a DDR factor enriched on the sex body (Royo et al., 2013) and also stained for BRG1, to demonstrate protein loss in mutant pachytene spermatocytes (Fig. S6A). Given that SCML2 physically associates with γH2Ax (Hasegawa et al., 2015; Luo et al., 2015), it may be possible that SCML2 localizes to autosomal sites harboring persistent γH2Ax in *Brg1^cKO^* spermatocytes (Fig 5A, panel insets). Surprisingly, subtle defects in SCML2 localization were also seen in *Brg1^Het^* pachytene spermatocytes, where it appeared more homogenously distributed genome-wide (Fig. S6B, panel insets).

**Figure 5.**
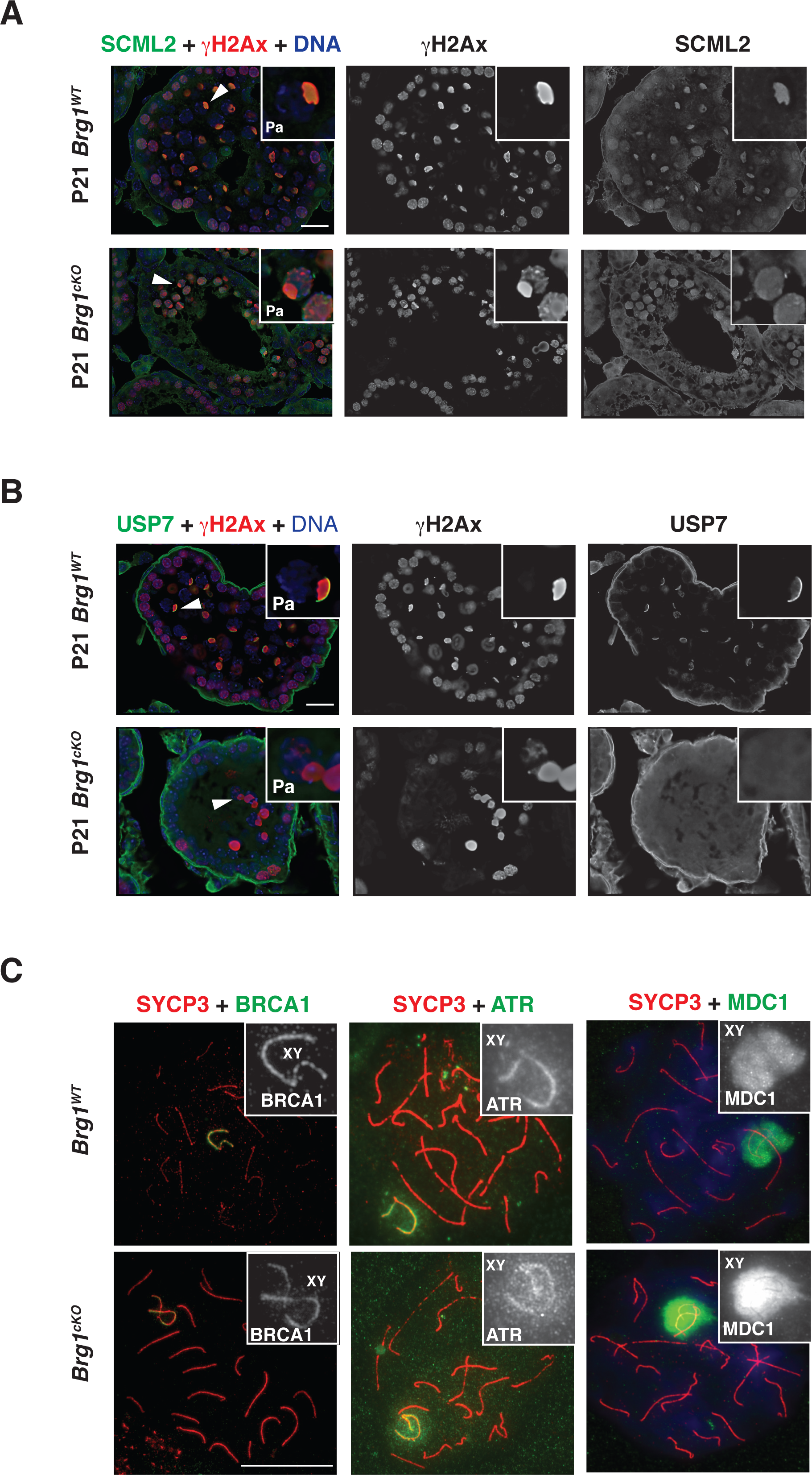
BRG1 influences the localization of SCML2 to the sex body. P21 *Brg1^WT^* and *Brg1^cKO^* testes cryosections (63x objective, Scale bar: 20 μm), co-stained for γH2Ax (red) and (A) SCML2 (green), (B) USP7 (green). DNA (blue) stained with DAPI. Arrowheads label the sex body. Panel insets show representative pachytene (Pa) spermatocytes. (C) *Brg1^WT^* and *Brg1^cKO^* pachytene spermatocytes spreads (100x objective, Scale bar: 20 μm), co-stained for SYCP3 (red) and left: BRCA1(green), middle: ATR (green), right: MDC1 (green). Panel insets highlight sex chromosomes.

We next examined SCML2 localization in *Brg1^WT^* and *Brg1^cKO^* meiotic spreads, co-stained with a synaptonemal complex marker, SYCP3, to visualize chromosomes. Mutant meiotic spreads were obtained from *Brg1^cKO^* testes generated using two independent germ line specific CRE’s, namely the *Mvh-cre* and *Stra8-cre*. Similar to the cryosections, SCML2 association with the sex chromosomes was perturbed in *Brg1^cKO^* pachytene spreads (Fig. S7). However, the phenotype appeared less severe in *Brg1^cKO^* pachytene spermatocytes generated with *Stra8-cre*, relative to *Mvh-cre* (Fig. S7). Such differences might be a consequence of the distinct temporal activity of each CRE (see materials and methods) (Gallardo et al., 2007; Sadate-Ngatchou et al., 2008).

In addition to SCML2 we also monitored the localization of USP7 in pachytene spermatocytes obtained from *Brg1^WT^* and *Brg1^cKO^* testes. Even though BRG1 does not directly interact with USP7 (Fig. S4C), we posited that the mis-localization of SCML2 in *Brg1^cKO^* pachytene spermatocytes might affect USP7 enrichment on the sex body. In fact this seems to be the case in *Brg1^cKO^* pachytene spermatocytes immunofluorescently stained for USP7 and γH2Ax (Fig. 5B).

From previous studies it is clear that the mis localization of SCML2 does not affect processes, critical to the initiation of meiotic sex chromosome inactivation (MSCI) (Hasegawa et al., 2015). Since the loss of BRG1 has been previously reported to influence MSCI (Wang et al., 2012), we examined its impact on the localization of known MSCI factors such as BRCA1, ATR and MDC1 to the sex body (Ichijima et al., 2011; Turner et al., 2004). By immunofluorescence, BRCA1, ATR and MDC1 appeared stably associated with the sex body in *Brg1^cKO^* pachytene spermatocytes (Fig. 5C). Thus BRG1 like SCML2 does not affect the initiation of MSCI. Interestingly, it was previously reported that the loss of MDC1 abrogates SCML2 recruitment to the sex chromosomes (Hasegawa et al., 2015). Therefore, the stable association of MDC1 with the sex body in *Brg1^cKO^* spermatocytes (Fig. 5C), suggests that SWI/SNF may function downstream MDC1 in the recruitment of SCML2 to the sex body.

### BRG1 influences the abundance of SCML2 associated histone modifications

Since SCML2 is known to regulate both H2AK119ub1 and H3K27ac during meiosis (Adams et al., 2018; Hasegawa et al., 2015), we investigated whether they are also affected by BRG1. We monitored the abundance of H2AK119ub1 and H3K27ac in acid extracts obtained from P12 *Brg1^WT^*, *Brg1^Het^* and *Brg1^cKO^* spermatogenic nuclei (Fig. 6A, top panel). We also examined the abundance of H3K4me3 and H3K27me3 which are associated with BRG1 target promoters (Fig. 6A, bottom panel). While neither H3K4me3 nor H3K27me3 levels were perturbed, both H2AK119ub1 and H3K27ac were dramatically elevated in P12 *Brg1^Het^* and *Brg1^cKO^*, relative to the *Brg1^WT^* spermatogenic cells (Fig. 6A, top panel). Interestingly, these epigenetic perturbations were undetectable at P8 and P10, which coincide with the initiation of meiosis and pairing (Leptonema to zygonema) (Fig. S8A). Thus BRG1 suppresses H2AK119ub1 and H3K27ac prior to the onset of pachynema. The dose dependency of such epigenetic regulation, prompted us to verify whether it was a consequence of the *Brg1* hypomorph or due to the mere presence of the *Mvh-cre*. The former scenario appears true, as evidenced by lack of any difference in the levels of H2AK119ub1, between males with the CRE and their littermate control (Fig. S8B). Another cause for concern was that, the H2AK119ub1 antibody (clone E6C5) used in this study was recently reported to recognize non-histone epitopes (Hasegawa et al., 2015). Hence, we validated the specificity of clone E6C5 by performing western blots on acid-extracted histones obtained from RNF2 (PRC1 E3-ubiquitiun ligase) knock out (*Rnf2^KO^*) embryonic stem (ES) cells, engineered using CRISPR-CAS9. The near absence of H2AK119ub1 signal in the *Rnf2^KO^* relative to *Rnf2^WT^* ES cell histone extracts confirms the specificity of clone E6C5 (Fig. S8C). Furthermore, H2AK119ub1 is clearly detected in chromatin fractions obtained from P12 *Brg1^WT^*, *Brg1^Het^* and *Brg1^cKO^* testes (Fig. S8D). Thus the changes in H2AK119ub1 reported here are genuine.

**Figure 6.**
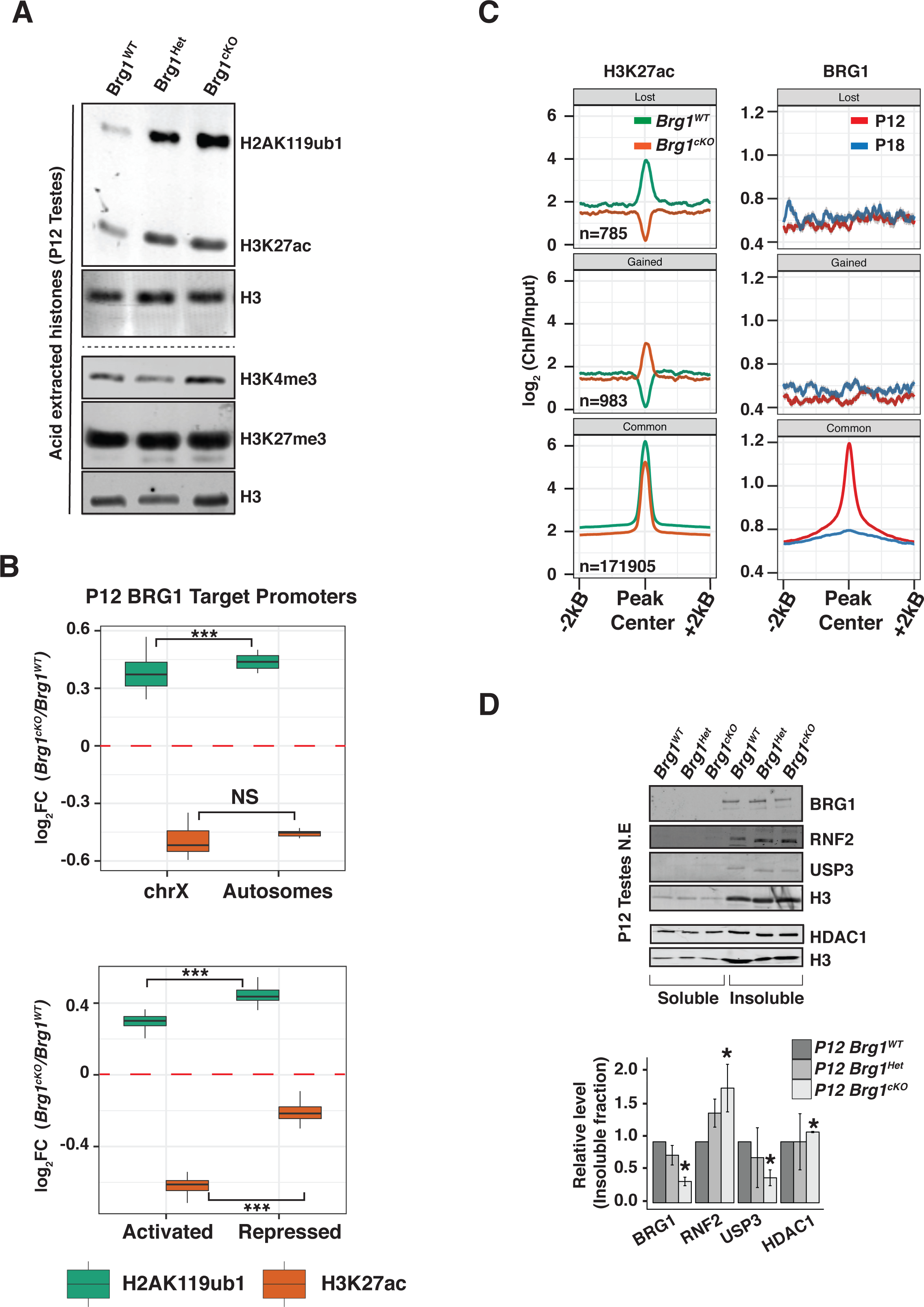
BRG1 regulates H2AK119ub1 and H3K27ac during spermatogenesis. (A) Western blots analysis of H2AK119ub1, H3K27ac, H3K4me3 and H3K27me3 in acid-extracts from P12 *Brg^WT^*, *Brg^Het^* and *Brg1^cKO^* testes. Loading control: Histone H3. (B) Log2 fold change (y-axis), in P12 H2AK119ub1 (green box) and H3K27ac (Orange box) at BRG1 occupied promoters (TSS± 0.5Kb, x-axis) categorized by chromosomal location (top panel) and transcriptional status (bottom panel). ***: p<0.001; NS: not significant, as calculated by Wilcoxon rank sum test. (C) H3K27ac (left) and BRG1 (right) enrichment at lost, gained and common H3K27ac peaks. (D) Western blot analysis (top panel) and quantification (bottom) of BRG1, RNF2, USP3 and HDAC1 in P12 *Brg^WT^*, *Brg^Het^* and *Brg1^cKO^* spermatogenic cells. Protein abundance are normalized to H3 and determined from at least two independent trials. Error bars represent the standard error of mean (SEM). *: p<0.05 (students t-test).

**Figure 7.**
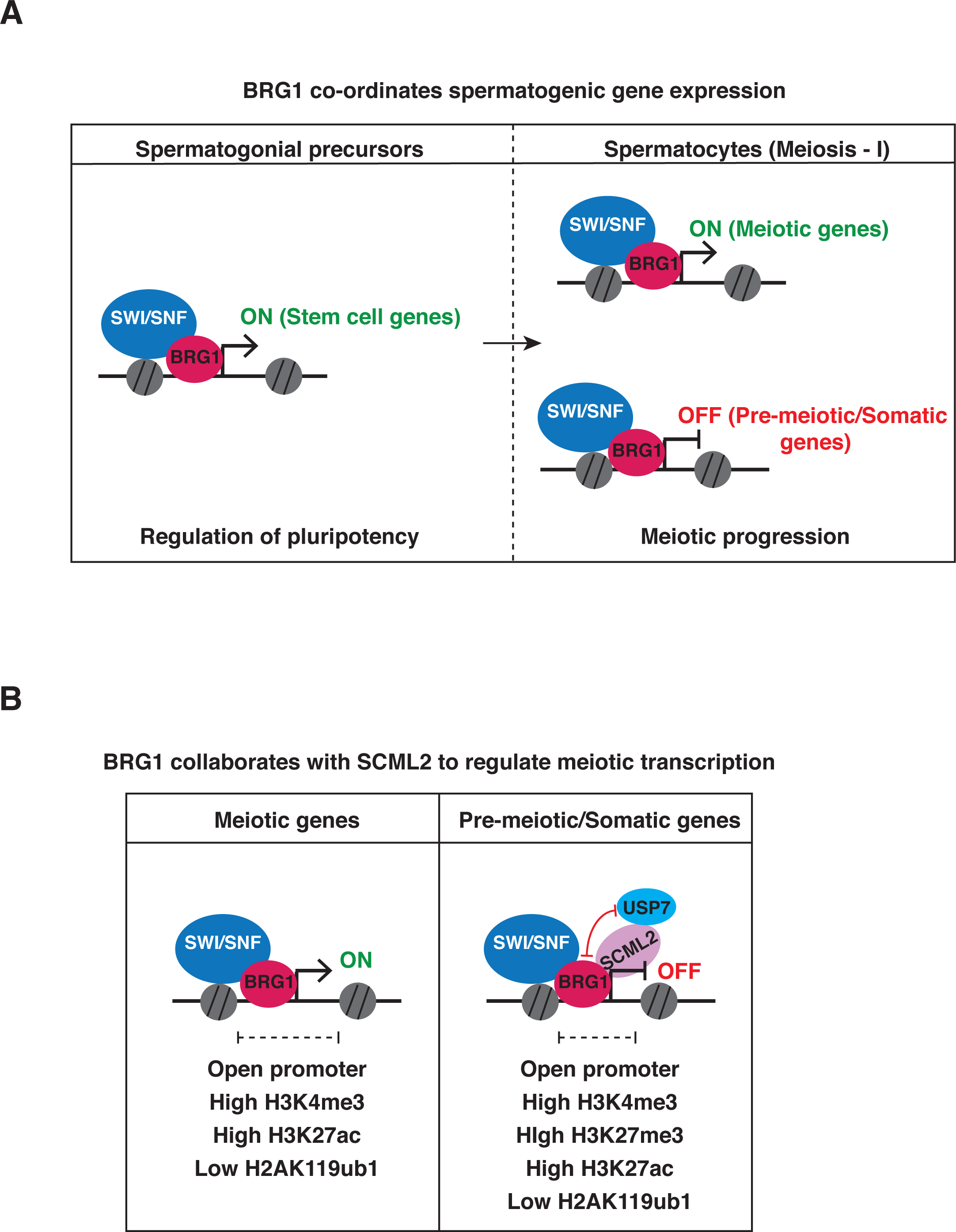
Model describing the role of SWI/SNF in spermatogenic gene regulation. (A) During spermatogenesis BRG1 activates genes essential for the maintenance of undifferentiated spermatogonia and ensures meiotic progression by activating meiotic genes and repressing pre-meiotic and somatic genes in spermatocytes (B) During meiosis, activated gene promoters display H3K4me3 while repressed gene promoters are bivalently modified (H3K4me3 and H3K27me3). BRG1 maintains promoter accessibility and suppresses H2AK119ub1 and enhances H3K27ac at target genes. We propose that BRG1 recruits SCML2 and its associated deubiquitinase, USP7 to epigenetically regulate cognate repressed targets.

### BRG1 suppresses H2AK119ub1 and enhances H3K27ac at target promoters

Based on the genome-wide increase in H2AK119ub1 and H3K27ac in *Brg1^cKO^* testes, we decided to survey the changes in these marks at promoters usually occupied by BRG1 (P12 peaks) by ChIP-seq in P12 *Brg1^WT^* and *Brg1^cKO^* spermatogenic cells. First, we categorized the BRG1 targets by their chromosomal location, the rationale being that both H2AK119ub1 and H3K27ac are known to be differentially regulated between the autosomes and sex chromosomes at pachynema (Adams et al., 2018). The BRG1 target promoters displayed contrasting changes in H2AK119ub1 and H3K27ac enrichment in the *Brg1^cKO^* spermatogenic cells. Irrespective of chromosomal location, target promoters displayed enhanced H2AK119ub1 combined with reduced H3K27ac in the *Brg1^cKO^* relative to *Brg1^WT^* spermatogenic cells (Fig. 6B, top panel). This mirrors the changes in H2AK119ub1 and H3K27ac that occur on the sex chromosomes in *Scml2^KO^* pachytene spermatocytes (Adams et al., 2018; Hasegawa et al., 2015). Thus, the regulation of H2AK119ub1 and H3K27ac at regions occupied by BRG1 may be influenced by SCML2, with which it is physically associated.

Next, we analyzed the changes in H2AK119ub1 and H3K27ac at promoters of BRG1 target genes that are differentially expressed. We hypothesized that these epigenetic modifications might dictate the transcriptional status of associated genes. Consistent with changes observed at all BRG1 occupied promoters (Fig. 6B, top panel), H2AK119ub1 was elevated while H3K27ac was reduced at promoters of mis-expressed target genes, irrespective of their transcriptional status in the *Brg1^cKO^* relative to *Brg1^WT^* spermatogenic cells (Fig. 6B, bottom panel). These epigenetic changes are typically associated with gene repression and thus only account for the silencing of mis-expressed genes in *Brg1^cKO^* spermatogenic cells. Therefore, the activation of mis-expressed target genes occurs by mechanisms independent of H2AK119ub1 or H3K27ac.

The loss of H3K27ac at promoters normally occupied by BRG1 in *Brg1^cKO^* spermatogenic cells is inconsistent with its genome-wide increase (Fig.6A; top panel, Fig. 6B). To determine whether other genomic regions acquire greater H3K27ac enrichment upon the loss of BRG1, we performed a differential peak calling analysis using the macs2 bdgdiff algorithm (Zhang et al., 2008). This analysis revealed several intronic and intergenic regions that lost, gained or maintained (common) H3K27ac peaks in *Brg1^cKO^* relative to the *Brg1^WT^* spermatogenic cells (Fig 6C; left panel, Fig. S9). Compared to the common peaks, regions that loose or gain H3K27ac are normally depleted of BRG1 (Fig. 6C, right panel). Thus while total H3K27ac levels are elevated, its local distribution appears more heterogeneous upon the loss of BRG1.

### BRG1 can also influence the epigenome in a SCML2 independent manner

While our data suggest that BRG1 regulates H2AK119ub1 and H3K27ac, through its interaction with SCML2, we cannot rule out the possibility that BRG1 might also influence these histone modifications by directly regulating the expression of cognate epigenetic modifiers. In fact, our ChIP-seq data identifies BRG1 peaks at promoters of epigenetic modifiers known to influence H2AK119ub1, H2BK120ub1 and H3K27ac (Fig. S10A). These include the H2A ubiquitin ligase, RNF2 (Wang et al., 2004), USP3 a H2A/H2B deubiquitinase (Nicassio et al., 2007) and the histone deacetylases, HDAC1 and HDAC2 (Gallinari et al., 2007) (Fig. S10A). The transcript abundances of all these targets are significantly altered in response to the loss of BRG1 (Fig.S10B).

In the case of the histone ubiquitin modifiers, *Rnf2* and *Usp3*, the former displayed an increase while the latter displayed a decrease in transcript abundance in *Brg1^cKO^* relative to the *Brg1^WT^* spermatogenic cells at P12 (Fig. S10B, row 1). This is consistent with significant increases in protein levels in *Brg1^cKO^* (RNF2: 88% increase, USP3:59% decrease) relative to the *Brg1^WT^* chromatin fractions prepared from testes (Fig. 6D, lane 4-6). Thus BRG1 may influence H2AK119ub1 by maintaining a balanced expression of RNF2 and USP3. Furthermore this represents an SCML2 independent mechanism of epigenetic regulation. Since USP3 has also been associated with mono ubiquitinylated H2B (Nicassio et al., 2007), we monitored the levels of mono ubiquitination of H2B lysine 120 (H2BK120ub1), a mark associated with gene activation and chromatin relaxation (Fierz et al., 2011; Minsky et al., 2008; Pavri et al., 2006). At P12, H2BK120ub1 appeared elevated genome-wide in *Brg1^cKO^*, relative to *Brg1^WT^* spermatogenic cells (Fig. S10C). The changes in the *Brg1^Het^* spermatogenic cells appear to be a product of unequal protein loading.

Similar to the histone ubiquitin modifiers, H3K27ac associated modifiers, *Hdac1* and *Hdac2* displayed reduced transcript abundance in the *Brg1^Het^* and *Brg1^cKO^* (Fig. S10B, row 2). At least in the case of *Hdac1*, we were unable to identify a corresponding depletion in protein levels by western blot (Fig. 6D, lane 1-6). Since HDAC1 has been shown to compensate for HDAC2 in various developmental scenarios (Ma et al., 2012; Montgomery et al., 2009; Yamaguchi et al., 2010) it is unlikely that the perturbation in H3K27ac levels in the *Brg1^cKO^* spermatogenic cells, occurs via mis-expression of HDAC1/2.

## Discussion

In this study we integrate genomic and proteomic approaches to show that SWI/SNF directed regulation of transcription, influences meiotic progression in males. In spermatogenic cells, BRG1 is overwhelmingly promoter associated which is distinct from what has been observed in other mammalian cell types and embryonic tissue (Alexander et al., 2015; Alver et al., 2017; Attanasio et al., 2014). This re-enforces the notion that cell or tissue specific associations influence SWI/SNF function during development (reviewed in Ho and Crabtree 2010). We propose a model in which the SWI/SNF ATPase, activates essential spermatogenic genes, while maintaining the repression of somatic genes. We show that BRG1 facilitates promoter accessibility of differentially regulated genes, which is consistent with the generally accepted mechanism of SWI/SNF (reviewed in Clapier et al., 2017). The activated genes play critical roles in the maintenance of undifferentiated spermatogonial cell populations and facilitate meiotic progression. Target stem cell factors include, *Zbtb16* and *Id4*. The latter specifically identifies spermatogonial stem cells (SSC) (Green et al., 2018; Oatley et al., 2011). The influence of BRG1 on SSC maintenance thus demonstrates a conserved role for SWI/SNF across various stem cell lineages (Ho et al., 2009; Lessard et al., 2007). The regulation of *Sycp2*, which is associated with synaptonemal complex assembly and homolog synapsis (Yang et al., 2006), potentially explains the incomplete synapsis and subsequent meiotic arrest seen in *Brg1^cKO^* spermatocytes (Kim et al., 2012). Thus the male sterility associated with *Brg1^cKO^* adults is a consequence of a shortage of germ line progenitors and essential meiotic factors.

The repression of the somatic transcriptome during meiosis is a feature shared with two other epigenetic regulators, namely, PRC2 and SCML2, a testes specific PRC1 factor (Hasegawa et al., 2015; Mu et al., 2014). Here we propose that BRG1 achieves the repression of its target genes by recruiting SCML2 activity. This is supported by the following observations; BRG1 (i) physically interacts with SCML2, (ii) is enriched at the promoters of genes regulated by SCML2 (iii) concordantly represses most genes commonly regulated by SCML2 and (iv) influences SCML2 localization during pachynema, based on a pronounced effect on sex body association. Interestingly, the transition to pachynema is accompanied by a reduction but not depletion of BRG1 from the X chromosome. This reduced level of BRG1 might be sufficient to recruit SCML2 to the sex body. In contrast to the repressed genes, BRG1 concordantly activates fewer SCML2 regulated genes. Most of these are associated with the regulation of mitotic spindle checkpoints, which in the context of meiosis occurs late into meiosis-I (Eaker et al., 2001).

In addition to these observations, BRG1 also appears to influence known SCML2 histone modifications, H2AK119ub1 and H3K27ac (Adams et al., 2018; Hasegawa et al., 2015). However unlike SCML2, BRG1 does not appear to differentially regulate autosomal and sex linked chromatin (Hasegawa et al., 2015). It is possible that such differential regulation does not manifest at pre pachytene stages. Alternatively, this may be indicative of distinct epigenetic outcomes associated with SCML2 dependent and independent mechanisms. The latter is demonstrated by the effect of BRG1 on the expression of potent H2AK119ub1 modifiers, such as RNF2 and USP3.

As counter-intuitive as it may seem, repressed BRG1 targets genes are associated with low H2AK119ub1 and higher H3K27ac levels. Hence, the repression of target genes probably occurs by alternative mechanisms. One possibility is that these targets are silenced by the formation of bivalent promoters (H3K4me3/H43K27me3),.typically associated with somatic developmental genes in the germ line (Hammoud et al., 2014; Lesch et al., 2013; Lesch et al., 2016). In fact 1/3^rd^ of BRG1 associated TSS’s appear bivalently modified from P12 to P17. Interestingly, SCML2 has been recently shown to influence the formation of bivalent domains during germ line development (Maezawa et al., 2018). Hence, such SCML2 activity might potentially govern the repression of BRG1 target genes.

In addition to H2AK119ub1, BRG1 also suppresses H2BK120ub1, which is associated with transcriptional activation and chromatin relaxation (Fierz et al., 2011; Minsky et al., 2008; Pavri et al., 2006). Hence, an increase in H2BK120ub1 might potentially de-repress genes in *Brg1^cKO^* spermatogenic cells. This might occur due to the mis-regulation of USP3 or mis-localization of USP7, two deubiquitinases, whose activities have been associated with H2BK120ub1 (Nicassio et al., 2007; Van Der Knaap et al., 2005). The evidence for USP7 is based on its function in *Drosophila* (Van Der Knaap et al., 2005). In contrast, mammalian USP7 appears to influence H2BK120ub1 in a catalytically independent manner (Lecona et al., 2015; Maertens et al., 2010, Huether et al., 2014).

In conclusion we reveal the transcriptional basis for the meiotic defects previously described in *Brg1^cKO^* males, and present a new paradigm for studying co-operation between SWI/SNF and PRC1 factors in the regulation of the epigenome. While recent studies have illustrated the propensity of BRG1 to evict canonical PRC1 members from chromatin in normal and oncogenic cell culture models (Kadoch et al., 2016; Stanton et al., 2016), the relationship between SWI/SNF and variant PRC1 factors remain unexplored.

## Supporting information

## Acknowledgements

We thank Magnuson lab members for helpful comments on manuscript preparation. This work was supported by National Institutes of Health grants R01GM101974 and U42OD010924.

## Data availability

Raw and processed data are deposited with GEO (GSE119179, for access, please enter token number: wjyxqkgozrgrpir).

## Materials and Methods

### Generation of *Brg1* conditional deletion and genotyping

*Brg1 floxed* (Sumi-Ichinose et al., 1997), *Mvh-Cre* (activated at ~ E15) (Gallardo et al., 2007) and *Stra8-Cre* (activated only in males at P3) (Sadate-Ngatchou et al., 2008) were maintained on a outbred background genetic background using CD-1 mice. *Brg1^fl/fl^* females were crossed to *Brg1^fl+^*;*Mvh-Cre^Tg/0^* males to obtain *Brg1^flΔ^*;*Mvh-Cre^Tg/0^* (*Brg1*^cKO^,), *Brg1^fl+^*;*Mvh-Cre^Tg/0^* (*Brg1*^Het^) and *Brg1^fl+^* (*Brg1*^WT^) littermate controls. Similar crosses were made to generate the *Stra8-Cre* induced conditional knockouts and littermate controls. Genotyping primers used in this study include: *Brg1^fl/+^* alleles - (F)- 5^’^ -CCTAGCCAAGGTAGCGTGTCCTCAT-3^’^; (R) 5^’^ -CCAGGACCACATACAAGGCCTTGTCT-3^’^, the excised allele (Δ)- reverse primer used above in combination with (F) 5^’^ -CTAACCGTGTATGTAGCCAGTTCTGCCT-3^’^, *Mvh-Cre* - (F) 5^’^ - CACGTGCAGCCGTTTAAGCCGCGT-3^’^, (R) 5^’^ -TTCCCATTCTAAACAACACCCTGAA-3^’^ and *Stra8-Cre* - (F) 5^’^ -GTGCAAGCTGAACAACAGGA-3^’^, (R) 5^’^ -AGGGACACAGCATTGGAGTC-3^’^. All animal work was carried out in accordance with approved IACUC protocols at the University of North Carolina at Chapel Hill.

### Disruption of Rnf2 by CRISPR-Cas9

The sequences of sgRNAs for Rnf2 are 5’CACCGTGTTTACATCGGTTTTGCG3’ and 5’AAACCGCAAAACCGATGTAAACAC3’. sgRNAs were cloned into pX330-U6-Chimeric_BB-CBh-hSpCas9 (42230, Addgene) using Golden Gate assembly cloning strategy (Bauer et al., 2015). Briefly, 5 x 10^4^ E14 ES cells were cultured on 60 mm dishes for one day and then transfected with plasmids expressing Cas9 and sgRNAs, along with a plasmid expressing PGK-PuroR (Addgene, cat. no. 31937) using FuGENE HD reagent (Promega) according to the manufacturer’s instructions. The cells were treated with 2 μg/ml puromycin for two days and recovered in normal culture medium until ES cell colonies grew up. *Ezh1* targeted colonies were verified by DNA sequencing.

### Immunofluorescence staining

Spermatocyte spreads were prepared as described (Peters et al., 1997) or by using a protocol adapted from (Wojtasz et al., 2009), that was used to generate “3D-preserved” spermatocytes. The latter entails a detergent spreading technique in which single-cell preparations obtained as described (Biswas 2018) were treated with 0.25% NP-40 for not more than 2 minutes. These spreads were used to view SYCP2 staining. All spermatocyte spreads were generated from 2- to 3-week-old mice. Prepared slides were either dried down and stored at −80° C or stored in phosphate buffered saline (PBS) at 4° C in the case of “3D-preserved” spermatocytes. Testis cryosections were prepared essentially as described before (Kim et al., 2012) with a few modifications. Briefly, juvenile/adult testes were fixed in 10% neutral buffered formalin at 4°C. After 20 minutes of incubation in NBF the tissue was sliced into half and then fixed up to 1 hour. Fixed tissue were washed 3 times in PBS at room temperature and then saturated through a sucrose series - 10% (30 min), 20% (30 min) and 30% (1 hr). The tissue were then incubated overnight in 30% sucrose/optimum cutting temperature (OCT) formulation at 4° C and subsequently embedded in OCT. Frozen sections were cut at 9 μm thickness. Antigen retrieval was performed for all antibodies used in this study. Briefly slides were incubated in boiling 10mM citric acid (pH6.0) for 10 minutes. Over this period, citrate buffer was replaced every 2 minutes with fresh boiling buffer and then allowed to cool down gradually for up to 20 minutes. Tissue section and spreads were washed in PBS followed by permeabilization in 0.1% Triton-X 100 and then blocked in Antibody dilution buffer (10% bovine serum albumin; 10% goat/donkey serum; 0.05% Triton-X 100) diluted 1:10 in PBS for 20 minutes before incubation with primary antibody overnight at 4°C. The following day, samples were again washed, permeabilized and blocked in ADB/PBS after which they were incubated for 1 hour with Alexa fluor-conjugated secondary antibodies. Immuno-stained slides were finally washed twice in PBS/0.32% photoflo (Kodak), once in H_2_O/ 0.32% photoflo and then counterstained with DAPI before mounting in Prolong Gold antifade medium (P-36931; Life Technologies). A list of all the primary antibodies used in this study are provided (Table S4). We used alexa fluor conjugated secondary antibodies at a dilution of 1:500. All imaging in this study was done on a Zeiss AxioImager-M2.

### Isolation of Spermatogonial stem cells and RNA extraction

Testes cell suspension was generated from 8-day-old *Brg1^WT^* (n=5), *Brg1^Het^* (n=1), *Brg1*^cKO^ (n=2) mice as described previously (Kubota and Brinster, 2008). Conditional deletions were generated using the *Mvh-Cre* transgene. Spermatogonial stem cells (SSC) were enriched using THY1^+^ microbeads (Miltenyi Biotec; 131-049-101) followed by their isolation on magnetic activated cell sorting (MACS) columns (Miltenyi Biotec; 131-090-312). RNA from SSC’s were isolated and purified using the PicoPure^TM^ RNA isolation kit (Life technologies; KIT0204).

### RT-PCR and qPCR

cDNA was synthesized using random primer mix (NEB) and ProtoScript^®^ II reverse transcriptase (NEB). Real time qPCR was performed using Sso Fast EvaGreen supermix (Bio-Rad) on a CFX96 thermocycler (Bio-Rad). A list of qRT-PCR primers used in this study are provided (Table S5).

### Isolation of spermatogenic cells

Spermatocyte enriched populations were isolated by methods described previously (Chang et al., 2011; Mu et al., 2014). Spermatocyte populations were enriched using percoll and cell strainers. Cell populations were then used for downstream applications such as RNA-seq, ATAC-seq, ChIP-seq and nuclear lysate preparations for immunoprecipitations.

### RNA-seq

RNA was extracted from spermatogenic cells obtained from 4 biological replicates of P12 *Brg1^WT^* and *Brg1*^cKO^ mice each. Conditional deletions were generated using the *Mvh-Cre* transgene. Cells were treated with the TRIzol reagent (Invitrogen) and total RNA was isolated and cleaned up using the Direct-zol RNA kit (Zymo). Sequencing libraries were prepared using a Kapa mRNA library kit as per manufacturer’s instruction and then sequenced on a Illumina Hiseq 4000 (50 bp reads, single end). Scml2 RNA-seq was previously published and is publically available under GEO accession number GSE55060 (Hasegawa et al., 2015).

### RNA-seq data analysis

Gene expression was quantified using kallisto (Bray et al., 2016). Transcript levels (counts) were summarized per gene using tximport (Soneson et al., 2016) and then imported to perform a differential analysis of gene expression using edgeR (Robinson et al., 2010). The mouse (mm9) gene/transcript annotations were retrieved using the ensembldb R package (https://github.com/jotsetung/ensembldb). Low abundance genes (counts per million <1 across 4 replicates) were filtered out in edgeR and significant differences in counts were called at a false discovery rate (FDR)≤ 0.05. The lists of differentially expressed genes are provided (Table S1). Anatomy ontology terms were curated from EMAPA (The Edinburgh Mouse Atlas Project) (Hayamizu et al., 2013) and the analysis was done on MouseMine (http://www.mousemine.org) (Motenko et al., 2015). Gene ontology analysis were performed using clusterProfiler (Yu et al., 2012).

### ChIP-seq

BRG1 ChIP was performed exactly as described previously (Raab et al., 2015). We performed the ChIP in duplicates on 4×10^7^ wild type (CD-1) spermatogenic cells obtained from P12 and P18 mice each. H2AK119ub1 and H3K27ac ChIPs were also performed in duplicate on spermatogenic cells obtained from P12 *Brg1^WT^* and *Brg1*^cKO^ mice using a method described previously for low chromatin inputs (Brind’Amour et al., 2015), with minor modifications (See supplemental materials for details). The BRG1 ChIP samples were sequenced on a Hiseq 2500 using v4 chemistry (50 bp reads, single end), whereas the H3K27ac and H2AK119ub1 ChIP samples were sequenced on an Illumina Hiseq 4000 (50 bp reads, single end). The antibodies used for ChIP are listed (table S4). H3K4me3, H3K27me3 ChIP-seq data were previously published and are publically available under GEO accession number GSE61902 (Mu et al., 2014). ChIP data analysis methods can be found in the supplemental materials.

### ATAC-seq

ATAC-seq was performed on spermatogenic cells isolated from 2 biological replicates of P12 *Brg1^WT^* and *Brg1*^cKO^ mice each. Only a single sample from 18 day old *Brg1^WT^* and *Brg1^Het^* mice were processed for ATAC-seq. Conditional deletions were generated using the *Mvh-Cre* transgene. ATAC-seq libraries were made as previously reported (Buenrostro et al., 2013), with the exception of a double-sided SPRI bead size selection step of each library, using 0.5x and 1x ratio of SPRI beads to obtain a library size range of ~150bp to ~2kb. All libraries were combined and sequenced on a single lane of Illumina Hiseq 2500 using v4 chemistry (50bp reads, single).

### Preparation of nuclear lysates

Nuclear extracts were prepared from spermatocyte enriched preparations as previously described (Chandler et al., 2012; Li et al., 1991) with minor modifications. These nuclear lysates were used for co-immunoprecipitations (co-IP) and for the mass spectrometric analysis of BRG1 immunopulldowns (supplemental materials).

### Identification of BRG1 interacting proteins by mass spectrometry

Proteins isolated from BRG1 IP and IP with a non-specific rabbit IgG were run approximately 2 cm below the bottom of the well of a precast SDS polyacrylamide gel (short gel). The short gel was stained with GelCode blue protein stain (ThermoFisher) and the lanes containing each sample were cut out and analyzed by mass spectrometry at the University of Massachusetts Medical School Mass spectrometry Core Facility (see supplemental materials).

### Preparation of sub cellular protein fractions

Cytosolic, nucleoplasmic (soluble) and chromatin (insoluble) fractions were prepared as described (Méndez and Stillman, 2000) from spermatogenic cells obtained from P12 and P21 *Brg1^WT^*, *Brg1^Het^* and *Brg1^cKO^* mice. Conditional deletions were generated using the *Mvh-Cre* transgene.

### Preparation of acid extracted histones

Histones were extracted from spermatogenic cells obtained from P12 *Brg1^WT^*, *Brg1^Het^* and *Brg1^cKO^* mice using acid extraction protocol described previously (Shechter et al., 2007). Conditional deletions were generated using the *Mvh-Cre* transgene.

### Western blotting

Protein samples were separated by polyacrylamide gel electrophoresis and then transferred to PVDF (Polyvinylidene Difluoride) membranes (Bio-Rad) using wet/ semi-dry transfer apparatus (Bio-Rad). Western blots were generated using the Li-COR Bioscience Odyssey fluorescent western blotting reagents. All the antibodies used in this study and their corresponding dilutions are listed (Table S4).

## Supplemental Information

### Supplemental tables

**Table S1.** List of genes differentially regulated by BRG1 at P12 (Excel sheet).

**Table S2.** List of regions displaying significant differences in chromatin accessibility in the *Brg1^cKO^* relative to *Brg1^WT^* (Excel sheet).

**Table S3.** List of BRG1 ChIP-seq peak calls and the genomic location of sites differentially bound by BRG1 between P12 and P18 testes (Excel sheet).

**Table S4.** List of antibodies used in this study

| Antibody | Vendor (catalog #) | Application |
| --- | --- | --- |
| Rabbit anti-BRG1 | Abcam (ab110641) | ChIP-seq (10 µg)<br>IF (1:500) |
| Rabbit anti- YH2Ax | Cell signaling (9718) | IF (1:1000) |
| Mouse anti- YH2Ax | Millipore (05-636) | IF (1:1000) |
| Mouse anti-SYCP3 | Abcam (ab97672) | IF (1:500) |
| Rabbit anti-SYCP3 | Abcam (ab154255) | IF (1:500)<br>WB (1:1000) |
| Rabbit anti-SYCP2 | Millipore (ABE2622) | IF (1:500)<br>WB (1:1000) |
| Goat anti- ATR (N-19) | Santa Cruz Biotech (sc-1887) | IF (1:50) |
| Mouse anti-SCML2 (1D12) | Developmental Studies Hybridoma Bank (PCRP-SCML2-1D12) | IF (1:10)<br>IP (10 µg) |
| Rat anti- RNAPII subunit B1 (phosphor CTD Ser-2) | Millipore (04-1571) | IF (1:200) |
| Mouse anti- H2AK119ub1 (E6C5) | Millipore (05-678) | ChIP-seq (10 µg)<br>WB (1:1000) |
| Rabbit anti- H3K27ac | Active Motif (39133) | ChIP-seq (5 µg)<br>WB (1:1000) |
| Rabbit anti- H2BK120ub1 | Cell signaling (5546) | WB (1:1000) |
| Mouse anti- H3K27me3 | Abcam (ab6002) | WB (1:1000) |
| Rabbit anti- H3K4me3 | Abcam (ab1012) | WB (1:1000) |
| Rabbit anti- H3 | Abcam (ab1791) | WB (1:5000) |
| Mouse anti- RNF2 | Santa Cruz Biotech (sc-1887) | WB (1:1000) |
| Mouse anti- USP3 | GeneTex (GTX128238) | WB (1:1000) |
| Mouse anti- USP7 | Santa Cruz Biotech (sc-1887) | IF (1:100)<br>IP (2 µg)<br>WB (1:1000) |
| Mouse anti- HDAC1 | Abcam (ab7028) | WB (1:1000) |
| Rabbit anti- PDGFRA | Cell signaling (3164) | WB (1:1000) |
| Mouse anti α TUBULIN | Developmental Studies Hybridoma Bank (12G10) | WB (1:1000) |
| Rabbit anti NCL (Nucleolin) | Bethyl (A300-711A) | WB (1:5000) |

**Table S5.**
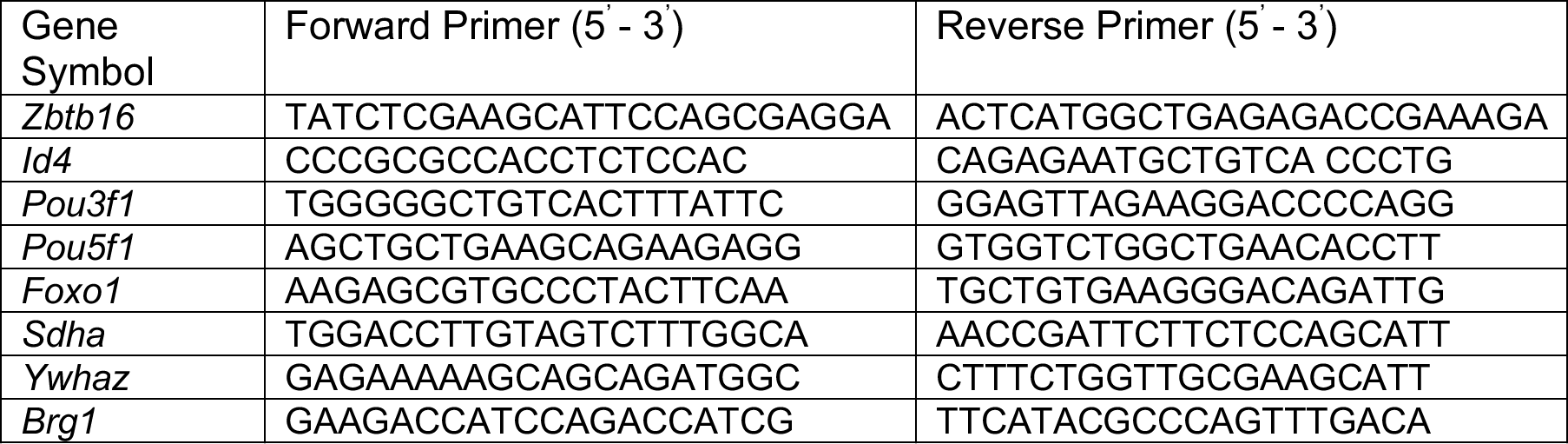
Sequences of qRT-PCR primers used in this study

### Supplemental figure legends

**Figure S1.**
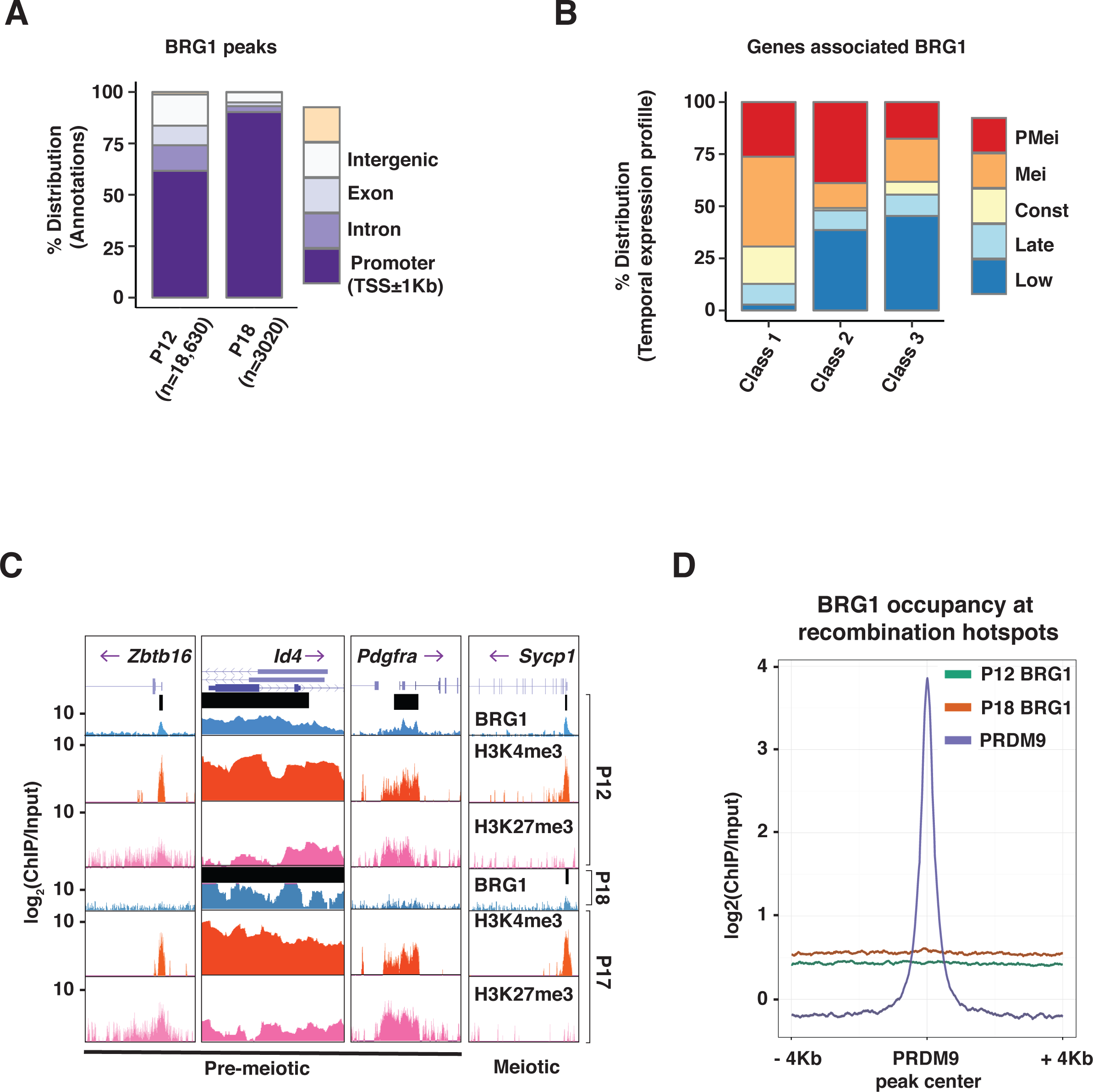
Features of BRG1 genomic associations. (A) Annotation of regions associated with P12 and P18 BRG1 peaks. Number (n) of P12 and P18 peaks are indicated within parenthesis. (B) Distribution of temporal expression profile of genes associated with class 1-3 TSS’s. PMei: pre-meiotic, Mei: Meiotic, Const: constant. (C) UCSC genome browser views depicting H3K4me3 and H3K27me3 peak associations with promoters of candidate BRG1 target genes. Thick black bars denote BRG1 Macs2 peak calls. (D) P12 (green line) and P18 (orange line) BRG1 enrichment across an 8 Kb window centered at PRDM9 peaks (purple line).

**Figure S2.**
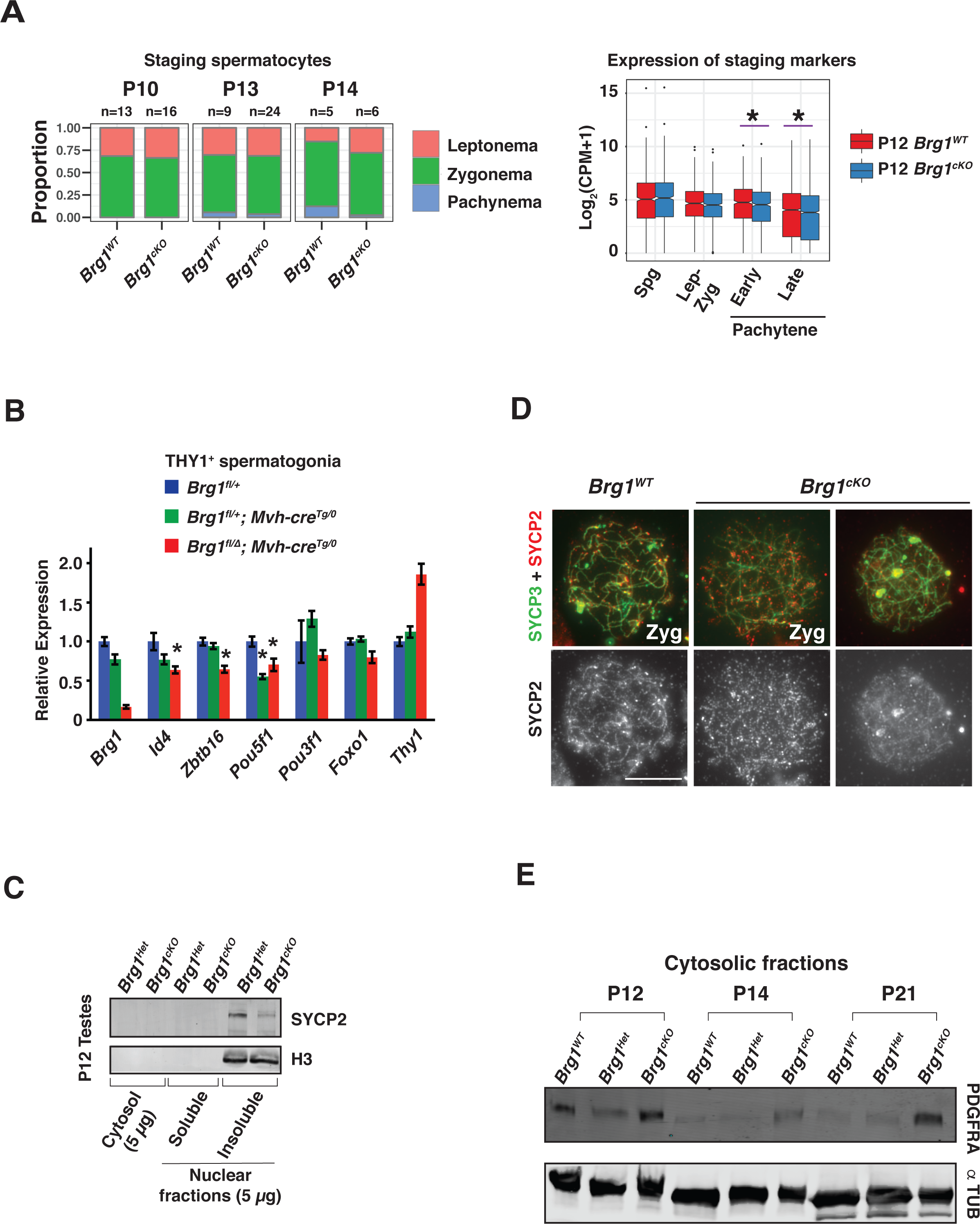
Transcriptional response to the loss of BRG1 in the male germ line. (A) Quantification of spermatocyte populations per tubule (left panel) staged by γH2Ax immunostaining of *Brg^WT^* and *Brg1^cKO^* cryosections obtained from P10, P13 and P14 testes. The total number of tubules analyzed (n) at each stage is indicated. Abundance (right panel) of spermatogonial and meiotic sub stage specific protein-coding genes (x-axis) between P12 *Brg^WT^* (red box) and *Brg1^cKO^* (blue boxes). Transcript abundance is expressed as the Log2 value of counts per million (CPM) added with a pseudo count (y–axis). (B) Quantitative RT-PCR analysis to determine the transcript abundance (y-axis) of candidate stem cell factors (x-axis) in *Brg^Het^* and *Brg1^cKO^* relative to *Brg^WT^* spermatogonia (THY1^+^). The transcript abundance of candidate factors was normalized to genes whose transcript levels are constantly expressed (*Sdha* and *Ywhaz*). * denotes a p-value < 0.05, calculated using an unpaired students t-test. (C) Western blot showing the abundance of SYCP2 in sub-cellular fractions obtained from P12 *Brg^Het^* and *Brg1^cKO^* spermatogenic cells. Histone-H3 was used as a nuclear loading control. (D) *Brg^WT^* and *Brg1^cKO^* zygotene spermatocytes immunofluorescently labeled for SYCP2 (red) and SYCP3 (green). Images were captured using a 100x objective Scale bar: 10 μm. (E) Western blot showing the abundance of PDGFRA in cytosolic fractions obtained from P12, P14, P21 *Brg^WT^*, *Brg^Het^* and *Brg1^cKO^* spermatogenic cells. α TUBULIN was used as a loading control.

**Figure S3.**
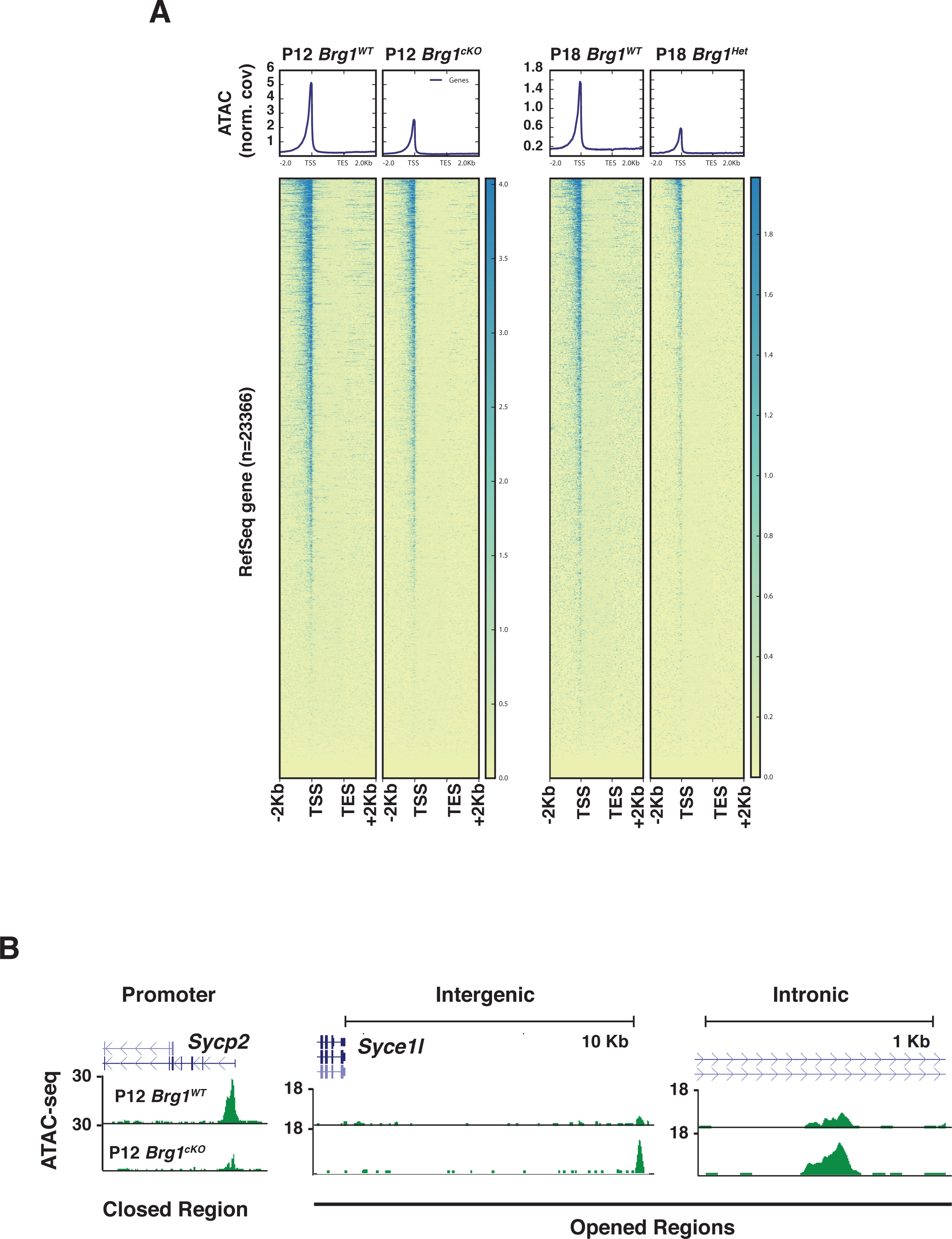
BRG1 directed changes in chromatin accessibility (A) Metaplots (top) and corresponding heatmaps (bottom) depicting the pairwise comparisons of normalized ATAC-seq signal at RefSeq genes ± 2Kb, between P12 *Brg1^WT^* and *Brg1^cKO^* spermatogenic cells, P18 *Brg1^WT^* and *Brg1^Het^* (*Brg1^fl/Δ^*) spermatogenic cells. TSS: Transcription start site, TES: Transcription end site. (B) UCSC browser view of candidate closed and opened regions.

**Figure S4.**
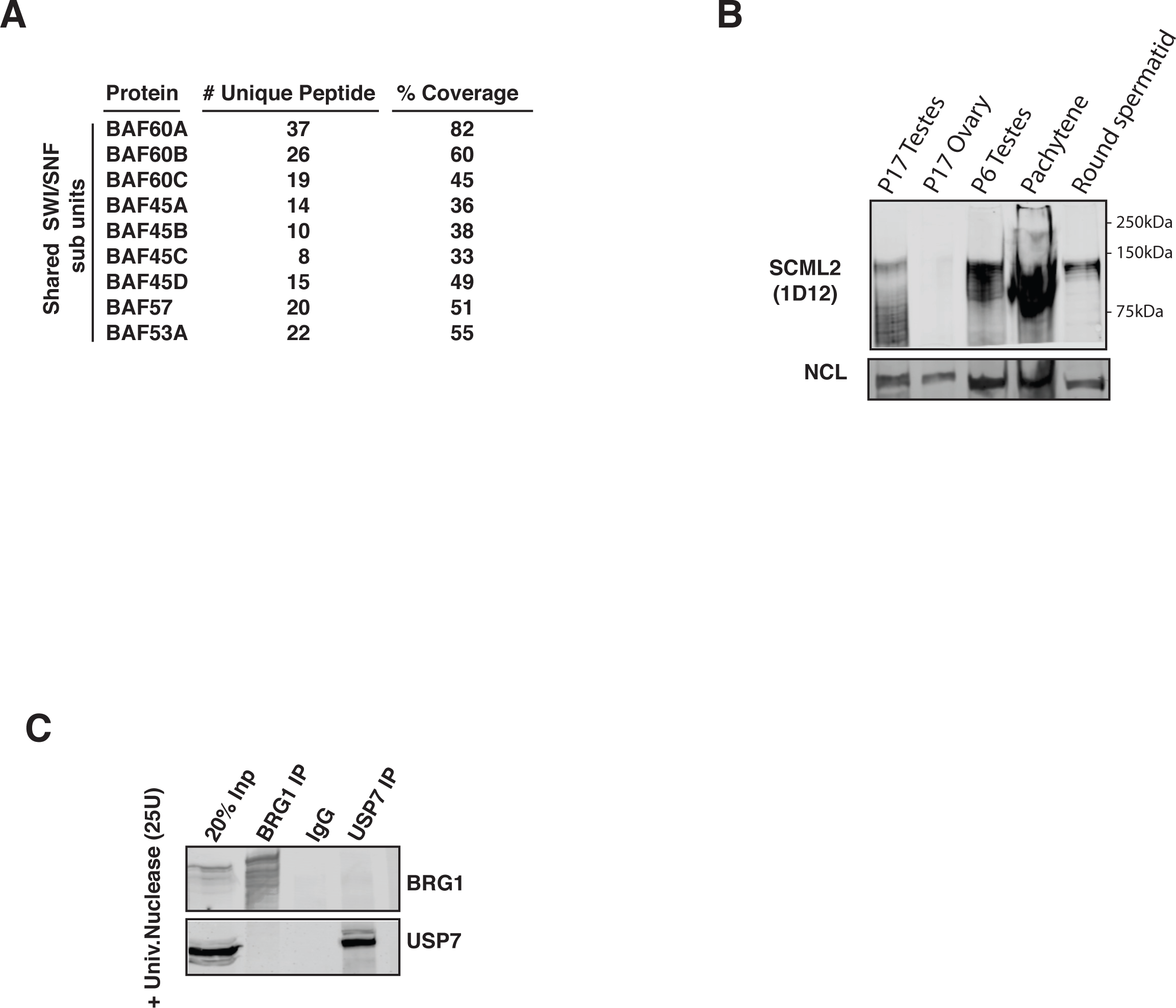
Summary and validation of BRG1 interactions. (A) List of known SWI/SNF subunits identified by IP-MS that are common to BAF and PBAF subcomplexes. (B) Validation of SCML2 antibody (1D12) specificity on nuclear extracts prepared from testes, ovaries and purified spermatogenic cells. Nucleolin (NCL) was used as a nuclear marker. (C) Co-immunoprecipitation of BRG1 and USP7. Nonspecific IgG was used as negative control. All lysates were treated with 25 U of universal nuclease (benzonase) prior to IP.

**Figure S5.**
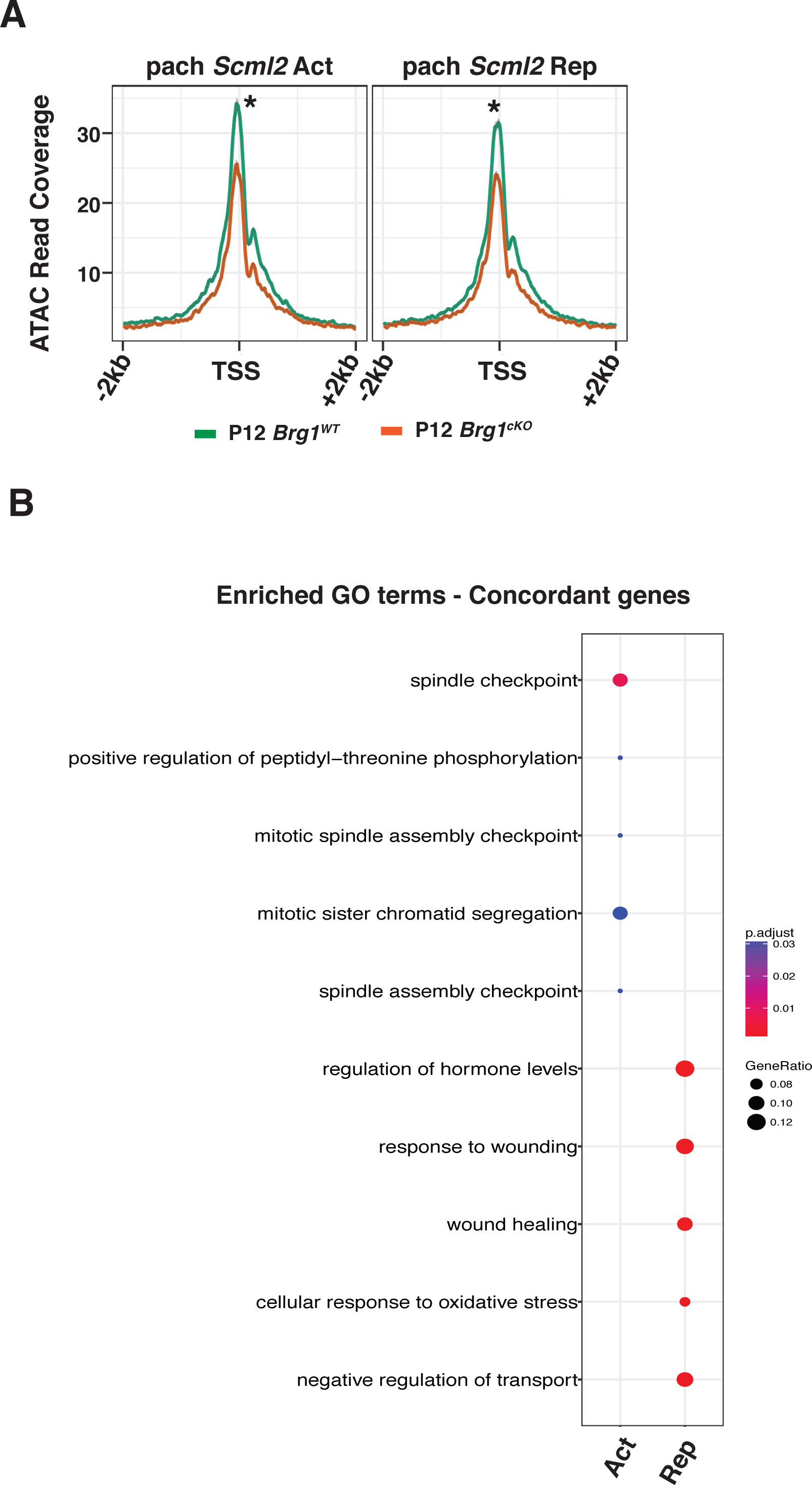
Features of BRG1 association with genes differentially regulated by SCML2. (A) ATAC seq coverage from P12 *Brg1^WT^* and *Brg1^cKO^* at TSS ± 4 Kb, associated with genes regulated by SCML2 in pachytene spermatocytes. (B) Enriched gene ontology terms associated with genes concordantly activated or repressed by BRG1 (P12) and SCML2 (pachynema).

**Figure S6.**
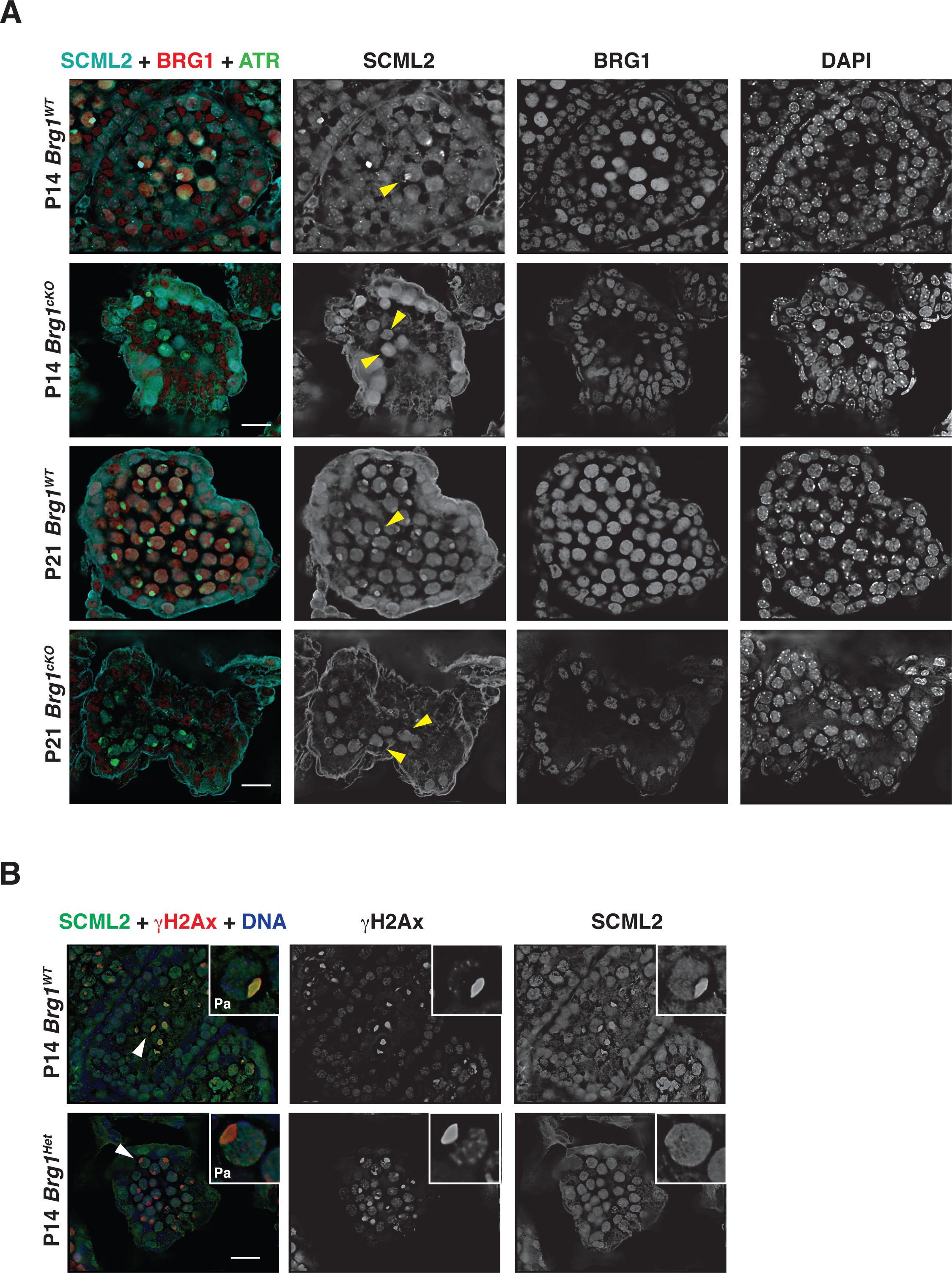
Validation of the role of BRG1 in the recruitment of SCML2 to the sex body. Cryosections prepared from (A) P14 and P21 *Brg1^WT^* and *Brg1^cKO^* testes, immunofluorescently labeled for SCML2 (cyan), BRG1(red) and ATR (green), (B) P14 *Brg1^WT^* and *Brg1^Het^* testes, immunofluorescently labeled for SCML2 (green) and γH2Ax (red), DNA was stained with DAPI. Arrowheads label pachytene spermatocytes with completely formed sex body. Images were captured using a 63x objective. Scale bar: 20 μm.

**Figure S7.**
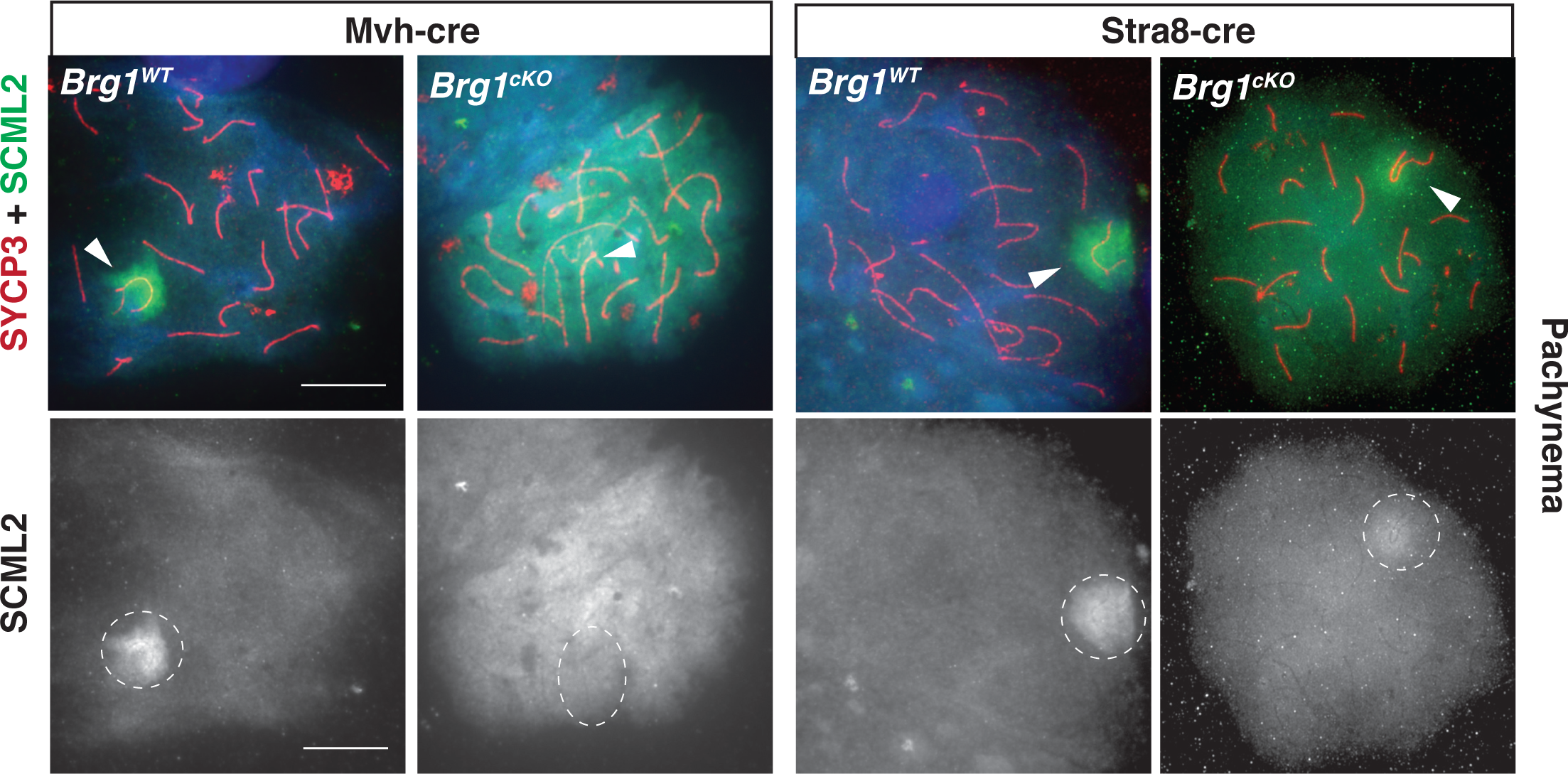
Analysis of SCML2 localization in meiotic spreads. Pachytene spermatocyte from *Mvh-Cre* (left panel) and *Stra8-Cre* (right panel) induced *Brg1^cKO^* and *Brg1^WT^* testes, immunofluorescently labeled for SCML2 (green) and SYCP3 (red). Arrowheads denote the sex chromosomes and dotted circle outlines the SCML2 signal around the sex body. Images were captured using a 100x objective. Scale bar: 20 μm.

**Figure S8.**
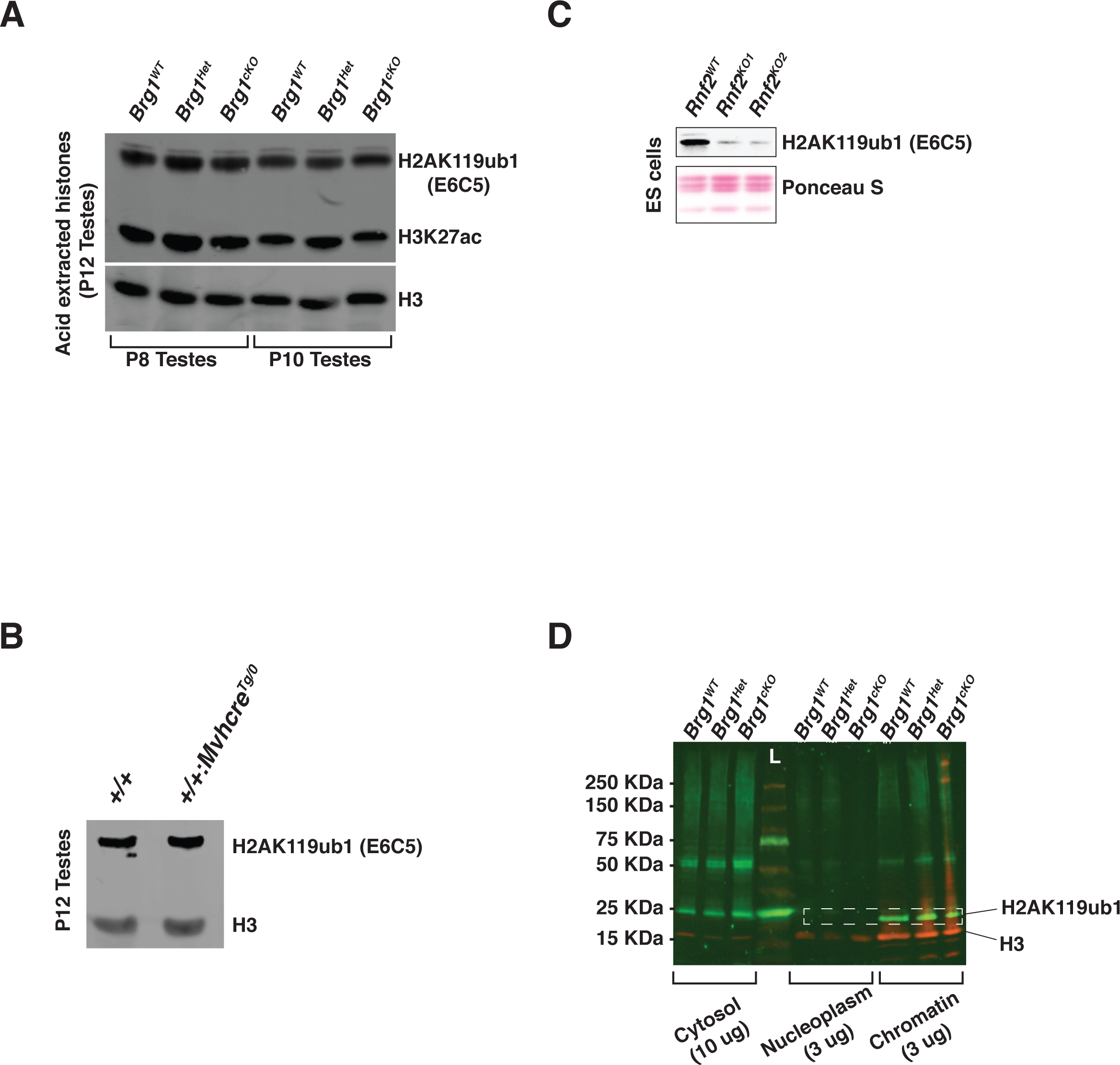
Temporal regulation of H2AK119ub1 in testes and validation of anti-H2AK119ub1 (clone E6C5). (A) Western blot depicting H2AK119ub1, H3K27ac abundance in acid extracted histones obtained from P8,P10 *Brg^WT^*, *Brg^Het^* and *Brg1^cKO^* testes. H3 was used as a loading control (B) Western blot showing H2AK119ub1 and H3 abundance in acid extracted histones obtained from testes of P12 wild type mice carrying a *Mvh-Cre* transgene and their littermate controls (*+/+*). (C) Validation of anti-H2AK119ub1 (clone E6C5) specificity by immunoblotting for total H2AK119ub1 in acid extracted histones obtained from *Rnf2^WT^* and two different *Rnf2^KO^* ES cell lines. Blots were stained with Ponceau S to show total histone levels. (D) Western blot using anti-H2AK119ub1 (clone E6C5), to show H2AK119ub1 (green) levels in sub-cellular fractions of spermatogenic cells obtained from P12 *Brg^WT^*, *Brg^Het^* and *Brg1^cKO^* testes. H3 (red) was used as nuclear loading control. Dotted rectangle labels the nuclear levels of H2AK119ub1 levels. L: ladder.

**Figure S9.**
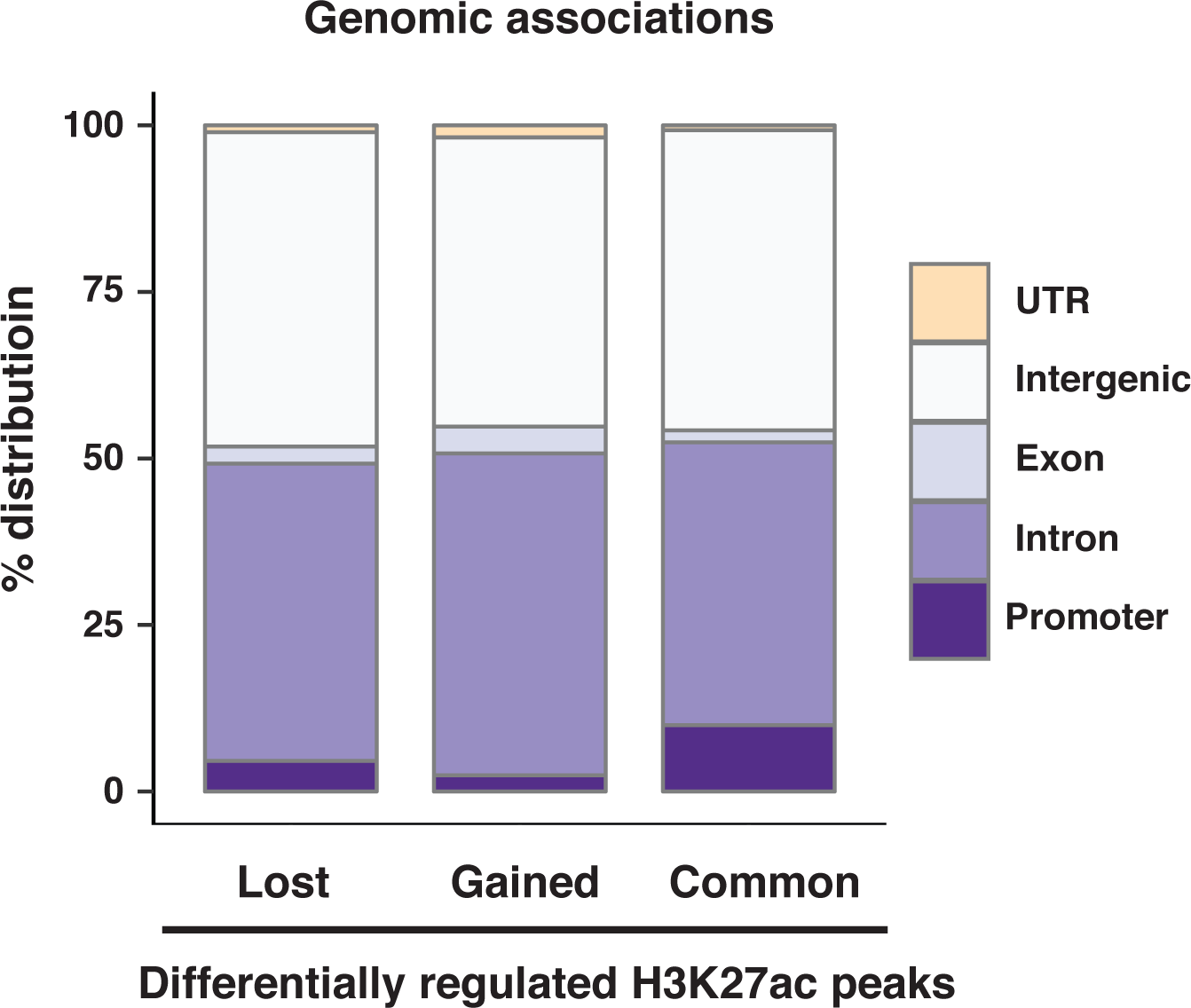
Genomic associations of lost, gained and common H3K27ac peaks.

**Figure S10.**
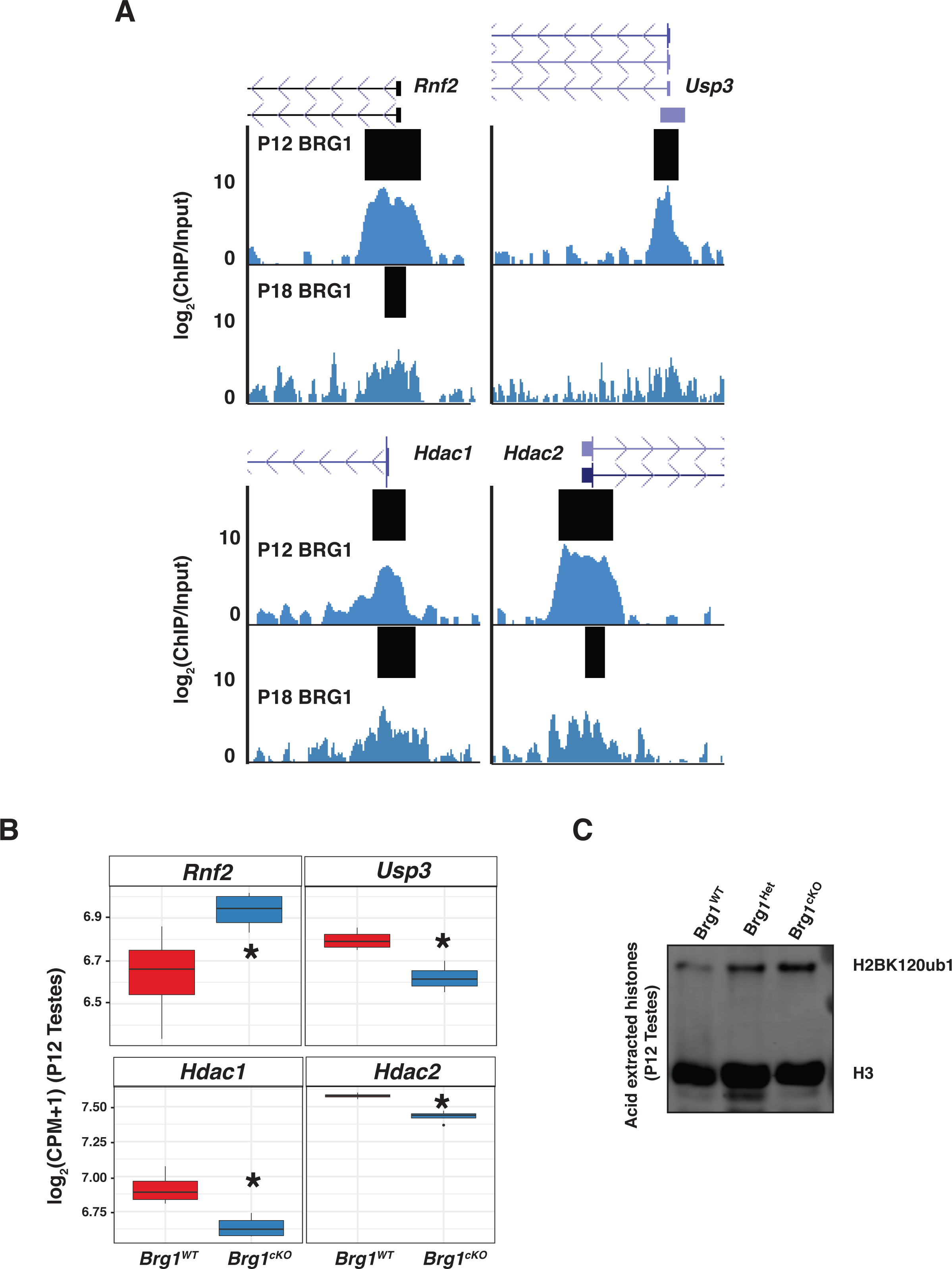
BRG1 regulates the expression of epigenetic modifiers of H2AK119ub1, H2BK120ub1 and H3K27ac. (A) UCSC browser view depicting BRG1 occupancy at *Rnf2*, *Usp3*, *Hdac1* and *Hdac2* promoters. Thick black bars label BRG1 peaks. (B) Transcript abundance (y-axis) of *Rnf2*, *Usp3*, *Hdac1* and *Hdac2* in P12 *Brg1^WT^* (red box) and *Brg1^cKO^* (blue box). Transcript abundance is expressed as the Log2 value of normalized abundance in counts per million (CPM) added with a pseudo count. (C) Western blot depicting H2BK120ub1 levels in acid extracted histones obtained from P12 *Brg^WT^*, *Brg^Het^* and *Brg1^cKO^* testes. H3 was used as a loading control.

## Supplemental Materials

### ChIP from low input chromatin

Spermatogenic cells obtained from 12-day-old *Brg1^WT^* and *Brg1*^cKO^ mice were fixed as described (Raab et al., 2015). Frozen pellets (10^6^ cells each) were thawed on ice and then resuspended in 50 μl of nuclear isolation buffer (Sigma NUC-101). For the H3K27ac ChIPs, 5mM sodium butyrate was added to the nuclear suspensions. Samples were mixed by pipetting the cell suspension 15-20 times. 50 μl of nuclear preparations were mixed with 10 μl of MNase digestion buffer [6x MNase buffer (NEB), 8.8 mM DTT, 12 gel units MNase (NEB)] and incubated at 37 C for 15 minutes. The digestion was stopped by adding 1/10^th^ volume of 100 μM EDTA (Ethylenediaminetetraacetic acid). 6.6 μl of a 1% Triton X-100/1% sodium deoxycholate solution was added to the digested chromatin and vortexed gently for 30 seconds after which the samples were held on ice until the next step. In the meantime the DNA content of each chromatin preparation was quantified from 5 ul of chromatin. Briefly, chromatin was mixed with 95 μl H_2_O and 100 μl 10% chelex-100 (Bio-Rad) and incubated at 95 C for 10 min. Following RNaseA digestion for 15 min at 37 C, ProteinaseK digestion for 1 hr at 56 C, the DNA was purified using ChIP-DNA clean and concentrator kit (Zymo) and quantified on a qubit. This way we ensured that for each pairwise comparison (wild type versus mutant ChIP), we began with an equal amount of input. We started with 145 ng and 285 ng of input DNA for each H3K27ac and H2AK119ub1 ChIP respectively. Digested chromatin was mixed with complete immnuoprecipitation (IP) buffer (20mM Tris-HCl pH 8.0, 2mM EDTA, 150 mM NaCl, 0.1% Triton X-100, 1x Protease inhibitor cocktail-PIC, 1mM Phenylmethanesulfonyl fluoride-PMSF) such that it comprised less than 25 % of the total volume. Chromatin was pre-cleared with 10 ul of magnetic Protein A (Bio-Rad) or G (Invitrogen/Dynalbeads) beads for 1 hour on a rotator at 4 C. After pre-clearing, 10% chromatin was set aside as input. Pre-cleared lysates were then mixed with antibodies to perform the ChIP. 5μg of rabbit anti-H3K27ac (Active Motif – 39133) conjugated to Protein A beads (Bio-Rad) and 10 μg of mouse anti-H2AK119ub1 IgM antibodies (Millipore E6C5, 05-678) were added to the lysates and left to bind chromatin by rotating the tubes overnight at 4 C. The following day anti-mouse IgM (Millipore, 12-488) conjugated to proteinA/G beads were added to the H2AK119ub1 ChIP samples and rotated at 4 C for 3 hours to capture anti-H2AK119ub1 bound chromatin. Chromatin bound to bead-antibody conjugates from all ChIP samples were isolated using a magnetic separator. Beads were then washed twice in low salt buffer followed by two washes in high salt buffer and eventually re-suspended in 100 μl of elution buffer (1%SDS/100mM NaHCO_3_) in a 1.5 ml eppendorf tube. DNA was eluted at 65 C on a shaking incubator (Eppendorf) at 800 rpm for 30 min. Eluate was separated from beads on a magnetic separator. 5 μl of 5M NaCl was added to eluate and incubated in 65 C water bath overnight to reverse crosslinks. ChIP DNA was digested with RNAseA for 30 min at 37 C followed by ProteinaseK digestion for 1 hr at 56 C and then purified using a ChIP-DNA clean and concentrator kit (Zymo) and quantified with a qubit. Libraries were prepared from ChIP and Input samples using the Kapa Hyperprep kit.

### ChIP-seq data analysis

Reads were aligned to mm9 using bowtie/bowtie2 (Langmead and Salzberg, 2012; Langmead et al., 2009). The resulting sam output files were converted to the BAM format using Samtools, version 1.6.0 (Li et al., 2009). The bam files were filtered to remove PCR duplicates using Picard tools, MarkDuplicates (http://broadinstitute.github.io/picard). The BAM files were converted to bigwig files for visualization on the UCSC browser (Kent 2002). The bigwig files were generated using DeepTools (Ramírez et al., 2016), bamCompare with a bin size of 10 bp, extending fragments to 150bp (nucleosome size), filtered for mm9 blacklisted regions and normalized to 1X depth of coverage. Each bigwig file represents the log_2_ ratio of ChIP to the corresponding Input sample. For comparison across samples the bigwig files were normalized to effective library size. Replicates were merged into a single bigwig using UCSC tools, bigWigMerge (http://hgdownload.soe.ucsc.edu/admin/exe/). Read coverage over regions of interest were generated from matrix files generated using DeepTools, computeMatrix following which metagene plots were made in R using ggplo2 (Wilkinson, 2011). Peaks from each sample were called using Macs2, version2.1.0, (Zhang et al., 2008) in broadpeak mode with--broad-cutoff 0.05 (FDR ≤ 0.05). Overlapping peaks between replicates were identified using bedtools, intersectBed and were used for subsequent analysis. A list of all BRG1 peak calls are provided (Table S3) Peaks were annotated using HOMER, peakannotate.pl (Heinz et al., 2010). Differential peak analysis was performed on bedgraph files using Macs2, bdgdiff run with default parameters.

### ATAC-seq data analysis

Reads were aligned to the mm9 genome using bowtie, with following paramters: -S –q –m 1 –p 2 –best –strata –chunkmbs 256 (Langmead et al., 2009). The outputed sam files were converted to BAM files using Samtools version 1.3.1. Next Bedtools version 2.25.0 (Quinlan and Hall, 2010), bamtobed was used to convert BAM output files to bed file format to do the subsequent steps. BAM files were converted to bigWig files for visualization on UCSC browser and to generate metagene plots as described above (see supplementary materials on ChIP-seq data analysis).

### Differential analysis of chromatin accessibility

For the differential analysis of chromatin accessibility, we adopted a method described for analyzing significant differences in counts between DNA hypersensitive sites identified by Dnase seq (Shibata et al., 2012). We compared differences in chromatin accessibility between 12 day old (P12) *Brg1^WT^* and *Brg1^cKO^* spermatogenic cells. Here the pairwise comparison was performed using edgeR, after obtaining the read counts from each replicate across defined 300 bp windows generated from a union set of the top 100,000 peaks (ranked by F-Seq). We called windows with significantly different counts from the pairwise comparison at a FDR ≤ 0.05. The significantly altered regions were annotated using HOMER, peakannotate.pl (Heinz et al., 2010). Regions with significant differences in open chromatin are provided (Table S2).

### Preparation of nuclear lysates for immunoprecipitation

Spermatogenic cells isolated from 2- to 3-week-old males were washed once in PBS and centrifuged at 600g for 5 min at 4 C. The resulting cell pellet was re-suspended in 20 PCV (packed cell volumes) of buffer A (10mM HEPES-KOH pH7.9, 1.5mM MgCl2, 10mM KCl, 0.1% NP-40, 0.5mM DTT, 0.5mM PMSF, 1x PIC, 0.5) and left on ice to swell for 10 min. Cells were centrifuged at 600g for 5 min at 4 C, resuspended in 2 PCV of buffer A and then homogenized using a dounce (type B). The homogenate containing nuclei were centrifuged at 700g for 10 min at 4 C. Nuclear preparations were washed once again in 10 PCV of buffer A and pelleted at 5000 rpm for 10 min at 4 C. Lysates were extracted from pelleted nuclei at least twice for 1 hour each with an equal volume of high salt buffer C (20mM HEPES-KOH pH7.9, 1.5mM MgCl2, 420mM NaCl, 10mM KCl, 25 % glycerol, 0.2mM EDTA, 0.5mM DTT, 0.5mM PMSF, 1x PIC) on a nutator at 4 C. The resulting nuclear lysates were mixed with 2.8x volume of buffer D (20mM HEPES-KOH pH7.9, 20 % glycerol, 0.2mM EDTA, 0.5mM DTT, 0.5mM PMSF, 1x PIC) following which they were clarified by spinning at 14,000 rpm for 10 min at 4 C, pooled together and snap-frozen in liquid nitrogen and stored at −80 C until further use.

### Co-Immunoprecipitation (Co-IP)

100 μl of magnetic protein A beads were initially washed twice in PBS for 5 min each at 4 C followed by one wash in PBS + 0.5% BSA for 10 min at 4 C. The antibodies (tableS3) were then mixed with beads resuspended in 1x volume of PBS + 0.5% BSA and allowed to conjugate for at least 2 hr at 4 C. After this, antibody-bead conjugates were washed once with 500 μl of PBS + 0.5% BSA, followed by two washes in IP buffer (20mM HEPES-KOH pH7.9, 0.15mM KCl, 10 % glycerol, 0.2mM EDTA, 0.5mM DTT, 0.5mM PMSF, 1x PIC) for 5 min each at 4 C. The antibody-coupled beads were stored in IP buffer until the nuclear lysates were processed for IP. Briefly, the lysates were thawed on ice and centrifuged at 14,000 rpm at 4 C to remove any precipitates. 500 μg of nuclear lysate was diluted in IP buffer to make up the volume to 1.3 ml and then incubated with unconjugated protein A beads to pre-clear the lysate. A fraction of lysate was set aside as input and the remaining was transferred to a tube containing the antibody coupled beads and left on a rotator overnight at 4 C. The following day the protein bound antibody-bead conjugates were separated using a magnetic separator and washed on a rotator with a series of buffers in the following order at 4 C for 5 min each: twice in IP buffer, twice in high salt wash buffer (20mM HEPES-KOH pH7.9, <u>300mM KCl</u>, 10 % glycerol, 0.2mM EDTA, 0.1 % Tween-20, 0.5mM DTT, 0.5mM PMSF, 1x PIC), twice in a low salt wash buffer (20mM HEPES-KOH pH7.9, <u>100mM KCl</u>, 10 % glycerol, 0.2mM EDTA, 0.1 % Tween-20, 0.5mM DTT, 0.5mM PMSF, 1x PIC), once in final wash buffer (20mM HEPES-KOH pH7.9, 60mM KCl, 10 % glycerol, 0.5mM DTT, 0.5mM PMSF, 1x PIC). The proteins were eluted by resuspending the beads in 2X Laemmli buffer and incubating them at 70 C for 10 min followed by boiling the samples at 95 C for 5 min.

### BRG1-IP and mass spectrometry

Prior to mass spectrometry the IPs were performed with anti-BRG1 antibody and non-specific IgG (table S4) as described above with a few modifications. After conjugation each antibody was crosslinked to protein A beads with BS^3^ (bis[sulfosuccinimidyl] suberate) crosslinking agent, prepared as per the manufacturer’s instructions (Thermo scientific, 21585). Crosslinking with BS^3^ significantly reduces IgG elution and improves signal to noise ratio in samples obtained by boiling beads in Laemlli buffer (Sousa et al., 2011). Briefly, antibody coupled beads were washed once in PBS for 5 min at 4 C, followed by two washes in conjugation buffer pH 7.9 (20 mM Sodium Phosphate, 0.15M NaCl). The antibody-coupled beads were resuspended in 250 ul of conjugation buffer containing 5mM BS^3^ in a tube and incubated on a rotator for 40 min at room temperature. Antibody crosslinked beads were separated and the crosslinker was quenched with an equal volume of 1M glycine (1x volume of beads). The crosslinked beads were then washed once in PBS at 4 C for 5 min, followed by incubation in 100 ul of 0.11M glycine (pH2.5) for 10 min at 4 C to remove any uncrosslinked antibody. The crosslinked beads were then washed thrice in PBS tween-20 (0.1%) + 0.5 % BSA, followed by another three washes in IP buffer before transferring them to a 15 ml falcon tube containing 4 mg of nuclear lysate made up in IP buffer up to a final volume of 10ml and left on rotator overnight at 4 C. The following day the beads were washed and proteins were eluted as described above. Samples were run on a short gel from which the regions containing the proteins were cut out and processed for mass spectrometry and peptide identification.

### In-gel Digestion

Gel slices were cut into 1×1 mm pieces and placed in 1.5 mL eppendorf tubes with 1 mL of water for 30 min. The water was then removed and 50 µL of 250 mM ammonium bicarbonate followed by10 µL of 45 mM 1, 4 dithiothreitol (DTT) were added prior to incubation at 50 °C for 30 min. The samples were cooled to room temperature and then alkylated with 10 µL of 100 mM iodoacetamide for 30 min. The gel pieces were washed 2x with 1 mL of water, removed and added 1 mL of 50 mM ammonium bicarbonate:acetonitrile (1:1) and allowed to incubate at room temperature for 1 hr. The solution was then removed and 200 µL of acetonitrile was added, removed, and the gel pieces dried by SpeedVac. Gel pieces were rehydrated in 70 µL of 2 ng/µL trypsin (Sigma) and 0.01% ProteaseMAX surfactant (Promega) in 50mM ammonium bicarbonate and incubated at 37 °C for 21hrs. Supernatant was removed, gel pieces added 100 µL of 80:20 (1% (v/v) formic acid in acetonitrile), combined with the former supernatant, and dried on a SpeedVac. Samples were reconsituted in 25 µL of 5% acetonitrile (0.1% (v/v) trifluroacetic acid) for LC-MS/MS analysis.

### LC-MS/MS

Tryptic peptides were dissolved in 0.1% trifluoroacetic acid and directly loaded at 4 μL/min for 7 minutes onto a custom-made trap column (100 μm I.D. fused silica with Kasil frit) containing 2 cm of 200Å, 5 μm Magic C18AQ particles (Michrom Bioresources). Peptides were then eluted onto a custom-made analytical column (75 μm I.D. fused silica) with gravity-pulled tip and packed with 25 cm 100Å, 5 μm Magic C18AQ particles (Michrom). Peptides were eluted with a linear gradient from 100% solvent A (0.1% (v/v) formic acid in water:0.1% formic acid in acetonitrile (95:05)) to 35% solvent B (0.1% (v/v) formic acid in acetonitrile) in 90 minutes at 300 nanoliters per minute using a Waters NanoAcquity UPLC system directly coupled to a Thermo Scientific Q Exactive hybrid mass spectrometer. Data were acquired using a data-dependent acquisition routine of acquiring one mass spectrum (*m/z* 300-1750) in the Orbitrap (resolution 70,000, 1e6 charges, 30 ms maximum fill time) followed by 10 tandem mass spectrometry scans (resolution 17,500, 1e5 charges, 110 ms maximum fill time, HCD collision energy 27 eV NCE). Dynamic exclusion was employed to maximize the number of peptide identifications and minimize data redundancy.

### Data Analysis

Raw data files were processed into peak lists using Proteome Discoverer (version 1.4; Thermo Scientific) and then searched against the Uniprot mouse database with Mascot (version 2.5; Matrix Science) using precursor mass tolerances of 10 ppm and fragment mass tolerances of 0.5 Da. Full tryptic specificity was specified considering up to 2 missed cleavages; variable modifications of acetylation (protein N-term), pyro-glutamination (N-term glutamine), and oxidation (methionine) were considered and fixed modifications of carbamidomethylation (cysteine) were considered. Mascot search results were loaded into Scaffold (Proteome Software) with threshold values of 80% for peptides (1.0% false-discovery rate) and 90% for proteins (2 peptide minimum) for final annotation.

